# Generation of a T cell receptor, cytokine and cell repertoire synovial fluid atlas to define commonalities and dissimilarities between arthritic diseases through systems immunology approaches

**DOI:** 10.1101/2025.01.10.632345

**Authors:** My K. Ha, Vincent Van Deuren, Carlos Alberto de Carvalho Fraga, An Hotterbeekx, Joke Dehoorne, Thomas Renson, Elke Geens, Nicolaas Aerts, Christiaan Heusdens, Maria Kuznetsova, Margot Van Mechelen, Michaël Vanden Bulcke, Ilse Hoffman, Ruth Wittoek, Eva De Wachter, Aurelie Andrzejewski, Esther Bartholomeus, Nick de Vrij, Arvid Suls, Eva Lion, Stijn Vanhee, Wim Adriaensen, Kerry Mullan, Helder Nakaya, Samir Kumar-Singh, Kris Laukens, Rik Joos, Pieter Meysman, Benson Ogunjimi

## Abstract

Although different chronic arthritic diseases are defined by clinical factors like gender, psoriasis and auto-antibodies, the biology of inflamed joints while comparing the different diseases remains neglected. Here, after curating an inflamed joint derived T-cell receptor (TCR) database, our new TRIASSIC tool identified 66303 significantly convergent TCR clonotypes. Clustering TCR clonotypes showed that synovial fluid convergence clusters (SFCCs) characterized HLA-B27+ mediated diseases (spondyloarthritis, SpA, and enthesitis-related juvenile idiopathic arthritis, JIA-ERA), Lyme arthritis and oligoarticular JIA. Single-cell transcriptomics and bulk proteomics showed upregulated interferon type I and II and TNF-α pathways in oJIA. Adult and juvenile psoriatic arthritis, (JIA-)PsA, was characterized by upregulated HSP expression in monocytes and TXNIP in T-cells. We discovered an abundance of CCL5 expressing CD8+ T-cells in SF from HLA-B27+ JIA-ERA and SpA patients. JIA-ERA patients showed upregulation of CD74 and LGALS1 in Th1 and Th17 cells and IGHV7-4.1 in B-cells. oJIA patients shared a TRBV28 RG-motif on CXCL13 producing helper T-cells. Rheumatoid arthritis and (JIA-)PsA patients carried EBV-reactive cytotoxic CD8+ T-cells. Annexin signalling was shown to be important in the intercellular communication for all arthritis groups. Collectively, our work showed that chronic arthritis is characterized by both disease-specific and broadly shared mechanisms.

## INTRODUCTION

Autoimmunity is the failure of an organism to recognize its own constituent parts as self, which results in an immune response against its own cells and tissues, causing autoimmune diseases. Chronic autoimmune arthritis, representing one major group of autoimmune diseases, can occur both in children and adults and can lead to phenotypically different disease groups. In adults we can distinguish at least three major disease groups: rheumatoid arthritis (RA), spondyloarthritis (SpA) and psoriatic arthritis (PsA). In children, juvenile idiopathic arthritis (JIA) consists of eight categories as defined by the International League of Associations for Rheumatology classification criteria: systemic JIA (sJIA), persistent oligo-articular JIA (oJIA), extended oligoarticular JIA (eoJIA), rheumatoid factor negative poly-articular JIA (pJIA), rheumatoid factor positive poly-articular JIA (RF^+^ JIA), enthesitis-related arthritis JIA (JIA-ERA), juvenile psoriatic arthritis (JIA-PsA) and undifferentiated JIA (uJIA) [1]. The JIA ILAR classification is highly debated and recently a new classification has been proposed [2,3], which might better reflect the underlying pathophysiology. Given the fact that therapy selection is typically based on (clinical) disease classification, it is of utmost importance to fully understand the different mechanisms leading to the different arthritic diseases. From this perspective, understanding the commonalities and dissimilarities in the synovial fluid and tissue between the different chronic arthritic diseases is crucial.

One understudied aspect is whether certain seemingly similar diseases in adults and children are driven by one shared overarching disease mechanism or rather distinct diseases. For example, adult SpA and JIA-ERA share clinical similarities such as sacroiliitis and enthesitis and are biologically characterised by a strong association with HLA-B27, which is a major histocompatibility complex (MHC) class I on the cell surface that is responsible for presenting antigenic peptides (derived from self and nonself antigens) to T-cells [4]. However, it is still unclear how and to what extent HLA-B27 contributes to the onset and pathogenesis of SpA and JIA-ERA [5]. There are some prevailing theories regarding the mechanism by which HLA-B27 exerts its effects. The molecular mimicry theory proposes that some structural features of HLA-B27, although unique amongst HLA molecules, may share molecular similarities with bacteria or viruses. An immune response to the infection with such bacteria or viruses would thus cross-react with HLA-B27, causing tissue damage and clinical symptoms [6]. The arthritogenic peptide theory is similar in hypothesizing that HLA-B27 binds to pathogen-derived peptides and triggers a CD8^+^ T cell response that is cross-reactive with HLA-B27/self-peptide pairs [6]. Specifically, the Salmonella outer membrane protein has been shown to elicit immune responses by SFMC from JIA-ERA and adult Reactive Arthritis and Undifferentiated SpA patients [7,8]. Other research has indicated that there exists an overlap between synovial fluid (SF) T cell receptors (TCRs) in SpA patients against self-antigens and microbial antigens [9]. This would imply that the same TCRs that allow T-cells to recognize microbes are the ones that cause inflammation in the joint. In the free heavy chains theory, a free cysteine (e.g., Cys67) promotes the formation of peptide-free, open conformation of HLA-B27 heavy chains (e.g., dimers), which participates in the pathogenic immune signalling by binding strongly to the leukocyte immunoglobin-like receptor family members (i.e., LILRB2 and LILRB5) in monocytes and osteoclasts, as well as the inhibitory killer cell receptor KIR3DL2 in natural killer cells, γδ T cells, activated CD4^+^ and CD8^+^ αβ T cells. The protein misfolding theory suggests that HLA-B27 misfolds in the endoplasmic reticulum (ER) due to several molecular factors (e.g., peptide binding specificity, folding kinetics, and the presence of unpaired cysteine residues) and thus causes ER stress which then leads to the inflammatory unfolded protein response [10], such as augmentation of IL-23 and activation of T_h_17 *in vitro* and in animal models [11,12]. Despite the prevalence of these theories, the exact mechanism by which HLA-B27 confers its adverse effects is still heavily debated. Although logic would dictate a biological similarity between juvenile and adult psoriatic arthritis, this remains to be studied. For example, when looking at the published TCR repertoires associated with adult PsA [13–16], we note that these have not been explicitly validated in JIA-PsA. In fact, in many studies the existence of different JIA subgroups was not considered.

Synovial inflammation in rheumatic and musculoskeletal diseases, particularly in RA, involves a complex interplay between immune and non-immune cell subsets. A key aspect of the immune response in seropositive RA is the adaptive immune response against citrullinated proteins, which includes both antigen-specific B cells and T cells [17]. Our understanding of rheumatoid arthritis (RA) is becoming increasingly comprehensive through single-cell atlases of both immune and non-immune populations derived from applying scRNA-seq to synovial tissue samples [18–21]. These investigations have revealed that the inflamed tissues of RA patients contain a diverse pool of cells in their various states (e.g., proliferation, migration, differentiation, etc) and that the presence and impact of these states can vary among different patient subgroups [20,21]. For T-cells, analysing TCRs offers valuable insights into developmental interaction between T-cell subsets and their expansion, capitalizing on the fact that each new T-cell produces a distinct TCR that is passed down to its progeny. Studies that examine the dynamics of TCRs across tissues or over time in RA patients have discovered shared T cell clones in various joints [22,23], clonal expansion of certain cell populations [24], the long-term persistence of expanded clones [25], and common gene rearrangements that may indicate shared antigenic affinity/recognition [22,26,27]. However, the reactivity between expanded TCRs from RA synovial CD4^+^ T cells and citrullinated peptides is yet to be certainly confirmed [28].

Thus, it is now necessary to conduct a thorough investigation and comparison of TCR repertoires in synovial fluid at inflamed sites across various autoimmune arthritic diseases to study the relationship of TCR clonal features with their functional roles, developmental pathways, and cell-cell interactions of specific lymphocyte phenotypes. From the TCR perspective, in the inflamed joints of JIA patients, a higher frequency of clonally expanded Treg cells was observed than conventional CD4^+^ T cells [29]. TCR profiling revealed the restriction and clonotypic expansion of Tregs in the blood and synovial fluid of JIA patients [30]. Between JIA patients, TCR overlap was noted between Treg cells in SF compared to Teff cells in SF [31]. Synovial fluid data from SpA patients showed some sharing of clonotype (clusters) [32]. Similarly, recent TCR research has indicated that CD8+ T cells capable of recognizing Epstein-Barr virus, cytomegalovirus, and influenza virus were present in the synovial fluid of adult rheumatoid arthritis (RA) patients [33]. Interestingly, SFMC-derived T-cell responses against human HSP60 are anti-inflammatory in oJIA, but not in pJIA or sJIA, and oJIA SFMC produce more IL-10 after stimulation with HSP60 [34,35].

This study leveraged the unique opportunity to simultaneously, thereby reducing interstudy biases, consider different types of autoimmune arthritis (including different JIA subtypes) at both cellular and molecular levels. Using single cell and bulk methods, we profiled the cell phenotype composition, gene expression, TCR repertoires and cytokine concentrations in the synovial fluid of patients having different autoimmune arthritis across different age groups. Our study is unique in the fact that we included all major adult and paediatric arthritic diseases. Our approach was centred around the “one group vs all-the-rest” principle where we compared biologicals from a baseline group with a “rest” group. Doing this we discovered novel insights regarding the underlying pathophysiology of the different chronic arthritic diseases and the potential pathogenic elicitors.

## RESULTS

### A large curated synovial fluid/tissue TCR database enables characterization and mapping of autoimmune arthritis associated convergent TCR signals

#### Extracting convergent TCR signals from synovial fluid and tissue

To study the antigen-specificity of the T-cells in the inflamed joints per disease group and compare across disease groups, we first compiled synovial fluid (SF) and tissue (ST) TCR-repertoire sequencing data from different studies describing autoimmune arthritic diseases into a database. This led to the curation of 242 SF/ST samples from 130 patients (Table 1). Starting from this TCR database, possible arthritis-relevant patterns were quantified in the TCR repertoire space. To overcome the inherent TCR repertoire diversity and sparsity, we developed a novel method called TRIASSIC that enabled us to analyze these heterogeneous TCR data and identify ‘convergent TCR clonotypes’. In brief, the TRIASSIC method identifies regions of the TCR space with high convergent recombination compared to control “background” repertoires (Figure S1a). This method is therefore based on the assumption that the disease-relevant TCRs are those that have more similar, independently generated, clonotypes in the arthritis group compared to the naive background. Through this approach, we successfully identified 66303 significantly convergent TCR clonotypes from the TCR database, which were clustered into 13516 **synovial fluid convergence clusters (SFCCs).** The cluster sizes varied, with 7013 of the SFCCs only containing one significant TCR (Figure S1b). In addition, most SFCCs (72.5%) occurred uniquely in one individual, indicating that they are private. This is expected as convergence and publicity are inversely correlated: public clusters are more unlikely to be highly convergent due to the prevalence of their TCRs across many repertoires. Inversely, those clusters that do not follow this typical pattern, and are therefore both public within a specific disease and convergent, potentially indicate disease-relevant patterns (Figure 1a). Indeed, such SFCCs capture several previously known arthritis-associated TCRs (Figure S1c).

**Figure 1.**
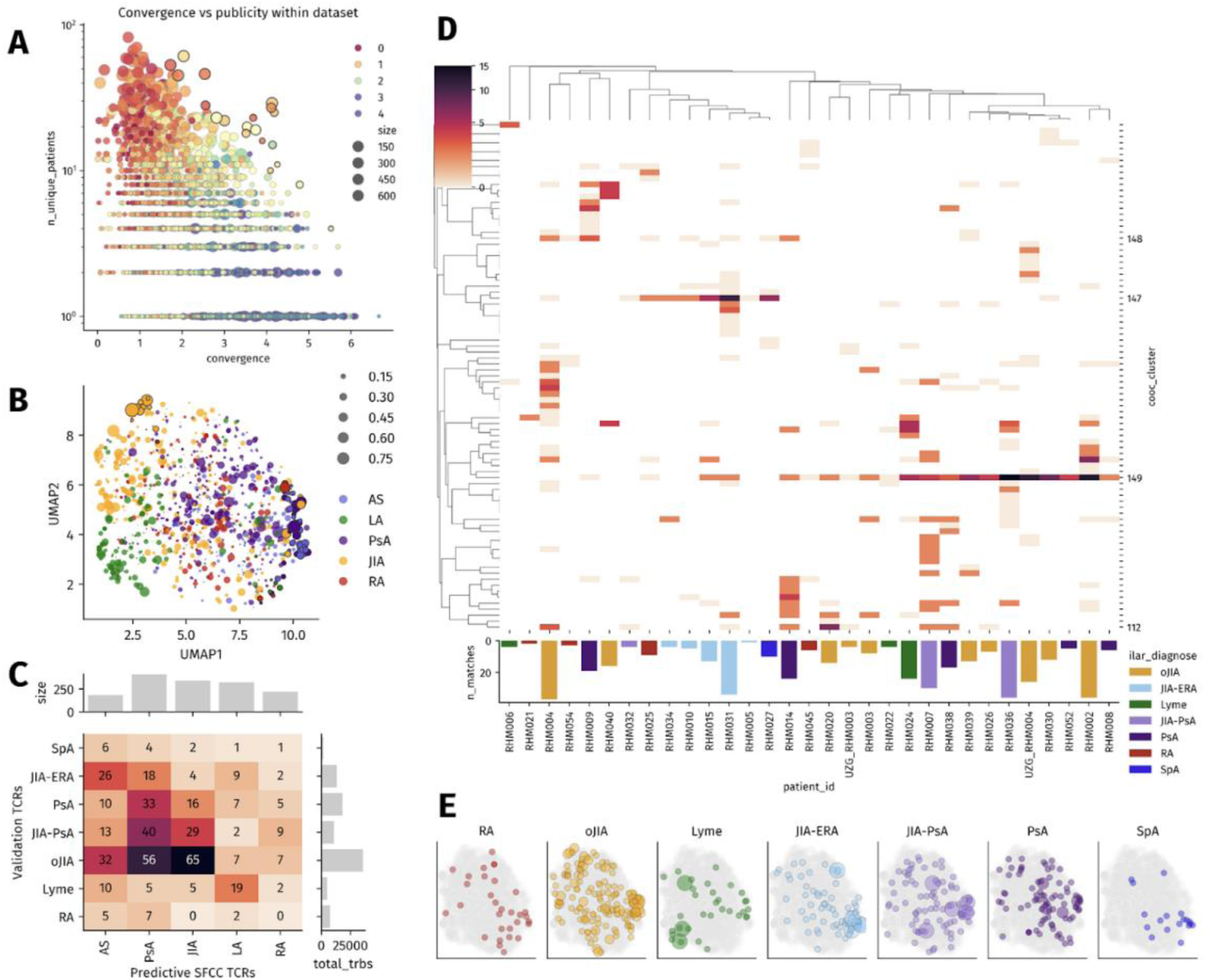
A. Highly convergent SFCCs (x-axis) are more likely to be private (y-axis), public TCR clusters are less likely to be convergent. **B**. Co-occurrence mapping reveals relationships among SFCCs using UMAP dimensionality reduction. Each dot represents a SFCC, and their spatial arrangement was determined as the cosine similarity between patient-level vectors (n=130 patients). Proximal points represent SFCCs with similar patient distribution patterns. Dot size represents the strength of association with arthritis disease subgroups determined by multiple logistic regression, while color indicates the diseases with the strongest association. Interestingly, several SFCC metaclusters emerged, comprising SFCCs associated with a specific disease. **C.** Cross-validation matrix showing the distribution of SFCC TCRs across disease groups. The x-axis represents the predictive SFCC TCRs associated with each disease group (AS, PsA, JIA, LA, and RA), while the y-axis represents the disease groups present in the validation dataset. Numbers in each cell indicate the number of TCRs from a SFCC in the validation dataset across all samples for a disease type. Bar plots on top and right show the total number of predictive SFCCs and sequenced validation TCRs, respectively. **D.** Hierarchical clustering of patients in the validation cohort based on SFCC metacluster abundances. The columns represent patients, with the x-axis indicating their disease group (by color), and the number of clonotypes sequenced for that patient (by height). The intensity in the heatmap (from light to dark) represents the abundance of each SFCC metacluster, with the most widely shared metaclusters 112, 149, 147, and 148 highlighted. **E.** Matching TCRs in the in-house validation data set for each disease group plotted on the same UMAP projection as in panel B, showing that disease-specific meta clusters remain consistent in the independent validation dataset.

**Table 1.** Clinical overview of patients.

| Category | Number | Percentage |
| --- | --- | --- |
| oJIA | 18 | 36% |
| JIA-ERA | 5 | 10% |
| JIA-PsA | 4 | 8% |
| PsA | 8 | 16% |
| RA | 7 | 14% |
| SpA | 4 | 8% |
| LA | 4 | 8% |
| HLA-B27 |  |  |
| • Positive | 8 | 16% |
| • Negative | 12 | 24% |
| • Undefined | 30 | 60% |
| ANA |  |  |
| • Positive | 23 | 46% |
| • Negative | 9 | 18% |
| • Undefined | 18 | 36% |
| RF |  |  |
| • *Positive* | 3 | 6% |
| • *Negative* | 4 | 8% |
| • *Undefined* | 43 | 86% |
| ACCP |  |  |
| • *Positive* | 3 | 6% |
| • *Negative* | 4 | 8% |
| • *Undefined* | 43 | 86% |

#### SFCCs can be validated in an independent dataset

To confirm that the found SFCCs were indeed disease-specific, we searched for the presence of these convergent signals across two independent datasets. First, we searched a database of control peripheral blood (PB) TCR repertoires comprising 4006 repertoires and over 325 million unique TCR clonotypes, compiled from seven large-scale TCR-seq studies (Table S2). Based on the reported metadata for the control PB repertoires, we considered this to be a set of healthy (non-arthritis) individuals. We found that SFCCs were indeed enriched in arthritis SF/ST compared to this control PB database. In addition, several SFCCs were entirely absent in the public PB despite spanning a large number of individuals, e.g. cluster 6886 (TRBV7-2 CATSREYGTsF[HY]NEQFF), which is present in 6 (4.7%) arthritis patients across different studies, yet in no controls. Secondly, we investigated the SFCC presence in an independent in-house validation SF-derived TCR dataset from 30 patients with various forms of autoimmune arthritis, containing both scTCR-seq and bulk TCR-seq data (Table 1).

#### SFCCs are arthritis disease-associated

Most samples within the arthritis database contain several of the identified SFCCs and their distribution is not random across individuals. Plotting the co-occurrence of SFCCs into a UMAP based on their presence in shared samples reveals distinct groups of SFCCs with similar occurrences in the database (Figure 1b). Next, we overlayed the putative HLA specificity of known public clonotypes in the SFCCs revealing several SFCC groups which have likely clustered together due to their targeting of the same MHC, and therefore possibly the same epitope (Figure S2). Next, we identified SFCCs that were the most predictive for specific chronic arthritis diseases using a L1-regularized multinomial logistic regression model. Similarly to the HLA-association, we note that SFCCs predictive for the same disease co-occur, hence clustering together in distinct regions of the SFCC landscape (Figure 1b). This was validated with our in-house SF: when overlayed on this graph, the most prominent disease-groupings were conserved (Figure 1c,e). These disease-associated groups included SFCCs with a known motif (e.g. TRBV9-associated TCRs in SpA), but also several previously undescribed motifs found by our novel TRIASSIC method. Despite these disease-associated signals, full unsupervised clustering of our in-house patient samples based on SFCC patterns could not fully separate different arthritis diseases (Figure 1d), even after filtering out non disease-associated SFCCs (Figure S3). Still, JIA-ERA and SpA patients are clearly delineated from other arthritic diseases by the presence of a distinct group of co-occurring SFCCs (metacluster 147), associated with HLA-B27. On the other hand, another prominent group of SFCCs (metacluster 149) was shared among patients with unspecified JIA, JIA-PsA, and PsA, and as these distinct diseases could not be homogeneously distinguished based on their TCR repertoire, thus indicating that they might share common epitope targets.

### In-depth transcriptomic characterization of SFCCs reveals shared, pathogenic T-cell signatures

Besides decoding antigen-specificity across diseases, the SFCC also allow the investigation of disease-specific T-cell transcriptomic signatures via our in-house cohort scGEX/TCR data. We selected three notable SFCC groups for a more detailed examination and functional characterisation using single cell gene expression and TCR data from our in-house cohort in the following sections.

#### EBV-reactive SFCCs drive inflammation across (JIA-)PsA and RA patients via cytotoxic CD8+ T-cells

Viral-reactive TCRs have been observed in the synovial fluid in multiple studies, thereby leading to the hypothesis that viruses might be involved in arthritis pathophysiology [36]. To investigate this, we annotated epitope specificity of the SFCCs using the ImmuneWatch™ DETECT tool (version 1.0) [37]. In total, 1.3% of the significantly convergent TCRs could be confidently annotated (IMW-DETECT score > 0.23), ultimately resolving the TCR specificity for 111 SFCCs. Notably, among the virus-specific TCRs annotations, those against Epstein-Barr virus (EBV) were the most common (36% of TCRs, 27/111 known-specificity SFCCs). Furthermore, TCRs from these clusters showed higher signs of convergent recombination when compared to those against other viruses (e.g. SARS-CoV-2) (Figure S4). Multiple of these EBV reactive SFCCs were found to be specifically associated with samples derived from PsA or RA patients. Furthermore, six of these EBV-reactive SFCCs were also present in the validation dataset, with every patient with (JIA-)PsA having at least one TCR from an EBV-reactive SFCC. On a phenotypic level, these six EBV-reactive SFCCs were CD8+ effector T-cells, significantly enriched in genes associated with cytotoxic functions (including GZMA, GZMB, GZMK, NKG7), and inflammatory cytokine production (including CCL5, CCL42L2) (see figure S). These findings suggest that EBV infection (acute, chronic or after reactivation) can induce cytotoxic T-cells that hold a causal role in some patients with autoimmune arthritis.

#### Broadly shared TRBV28 RG-motif SFCCs characterizes CXCL13 producing peripheral helper T-cells in oJIA

Unsupervised SFCC-based clustering of our in-house patient repertoires highlighted two distinct patient groups, driven by differential presence of groups of co-occurring SFCCs (SFCC meta-clusters). Within the most abundant SFCC meta-cluster (149), we noted a conserved TCR pattern consisting of a TRBV28 TCR with an RG motif in the hypervariable part of the CDR3 sequence, which is present in four different SFCCs. Within the curated TCR database, these SFCCs were primarily associated with JIA and PsA patients. The TCRs for these SFCCs were present, but uncommon in PB TCR repertoires of non-arthritis individuals compared to other SFCCs (Figure S5). All four clusters had matching TCRs in our in-house validation cohort, for a total of 59 matching unique TCR clonotypes, thereby allowing detailed phenotyping (Figure 2). Interestingly, oJIA patients had more TRBV28-RG SFCCs matches (51) than other groups, resulting in a significant enrichment (MWU p=3.716e-05, Figure 2). Nine out of ten oJIA patients had at least one hit, compared to 5/22 non-oJIA arthritis patients, indicating a conserved and unexpectedly broadly shared TCR signature amongst oJIA patients. Furthermore, in our single-cell GEX/TCR data, the SFCC TRBs paired with a strongly conserved TRAV8-3/TRAJ27 alpha chain, further indicating shared specificity (Figure S5b). According to IMW-DETECT, epitope specificity could not be annotated, indicating that they are rare or likely not associated with well-characterized viral epitopes. Phenotypically, these cells were an unconventional peripheral helper T-cell (Tph)-like population, marked by high CXCL13 production, ICOS, TNFRSF18, FOXP3 negativity, activation and exhaustion genes (HLA-DRB1, PDCD1, LAG3) (Figure S5e). We thus identified for the first time an oJIA-specific TCR signature that originates from unconventional Tph-like T-cells and could likely attract pathogenic B-cells through their production of the B-cell chemoattractant CXCL13.

**Figure 2.**
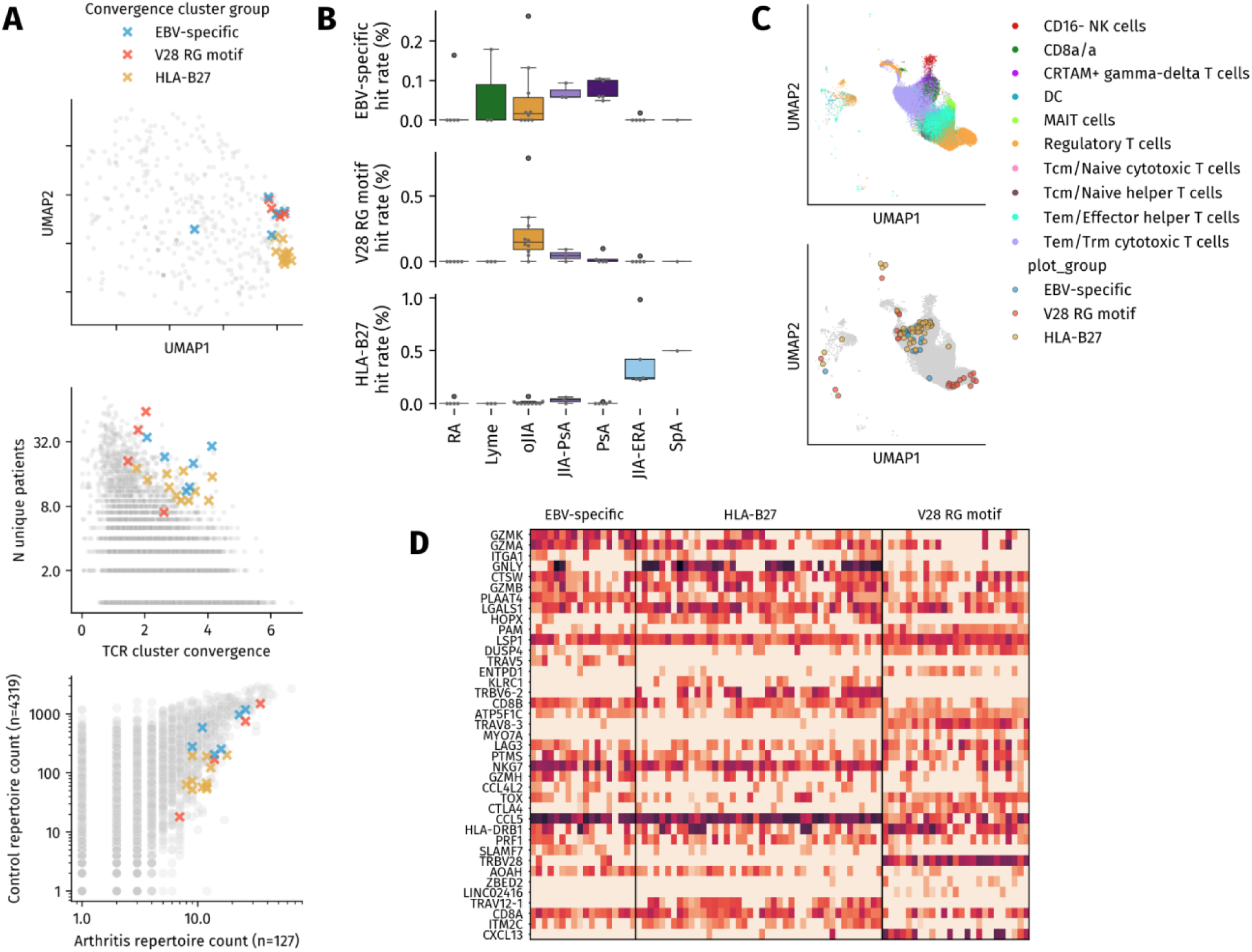
Phenotypic characterization of synovial T-cell clusters. **A.** Overview of features from selected SFCCs within the SF/ST TCR database. Top: UMAP representation visualization of SFCCs showing their position in the co-occurrence landscape. Middle: Publicity analysis of selected SFCCs showing the number of unique patients contributing TCRs (y-axis) versus TCR cluster convergence score (x-axis), demonstrating both high publicity and convergence. Bottom: selected SFCCs are enriched in arthritis vs control repertoires. **B.** Distribution of the SFCC groups across arthritis diseases in our in-house cohort. Each dot represents an individual patient and show the fraction of its SFCC-annotated TCRs clonotypes that belong to a specific SFCC group (y-axis). Plots are stratified by patient disease group (x-axis), with box plots representing their median and interquartile ranges. Statistical significance was assessed using Kruskal-Wallis test across all groups, followed by pairwise Mann-Whitney U tests for specific disease comparisons. Disease-specific enrichments include: oJIA vs non-oJIA for TRBV28 RG motif (p=3.716e-05), ERA/SpA vs others for HLA-B27-associated TCRs (p=2.588e-05), and LA vs others for LA-specific clusters (p=9.156e-03). **C.** T-cell phenotypes of SFCC-associated cells visualized using UMAP dimensionality reduction of single-cell gene expression data. The top panel shows T-cell subtype annotations based on Celltypist. Bottom panel highlights locations of cells belonging to one of the four investigated SFCC groups on the same UMAP coordinates. Notably, this plot highlights a conservation of T-cell phenotype within the same SFCC group, even across patients. **D.** Heatmap showing the expression of key differentially expressed genes within single T cells (rows) stratified by SFCC group. We performed differentially expression analysis for all T-cells from a single SFCC group (determined by their TCR), comparing to all other T-cells by t-test. The top 15 most significant DEGs per SFCC group were selected for display. The colour intensity indicates the mean expression level of a gene within a cell.

#### Multiple HLA-B27 associated SFCC denote SpA and PsA and originate from CCL5 expressing CD8+ T-cells

A second set of 20 co-occurring SFCC (meta-cluster 147) was distinctly associated with SpA and HLA-B27 positive PsA in our arthritis TCR database. Notably, 12 (60%) of the SFCCs from this group could also be annotated as HLA-B27-associated based on enrichment analyses from repertoires with known HLA status. Four of the clusters from this set of SFCCs are (variants) of the previously described TRBV9 CASS.GL[F/Y]STDTQYF TCR, which has been shown to play a pathogenic role in the development of SpA [38]. However, the remaining 16 SFCCs from this meta-cluster are strongly dissimilar and were not previously reported (e.g., TRBV7 CASS.RD.PYEQYF), indicating potential alternative causative motifs. In our in-house validation dataset, we see that TCRs matching to these SFCCs were significantly more prevalent in our (all HLA-B27+) JIA-ERA and SpA patients vs other arthritic diseases (MWU p=2.588e-05). Overlaying the single-cell data shows that T-cells from these SFCCs are phenotypically homogeneous CD8+ T-cells, with high CCL5 expression, possibly contributing to immune cell recruitment to the joint, and a strong cytotoxic signature, including upregulation of NKG7, GNLY, GZMB, and PRF1. Further HOPX expression suggests these are likely tissue-resident memory cells.

#### Pseudo-bulk single-cell derived differential gene expression and gene set enrichment analyses of synovial fluid shows disease-specific signatures

We previously illustrated how single cell gene expression of synovial fluid inflammatory cells can uplift the interpretation of our TCR data modeling. To investigate this in more detail (including other immune cell types beyond T-cells with specific receptor motifs), we analysed our synovial fluid derived single cell gene expression data between the different arthritic diseases in a cohort of 27 patients (Figure 3A). First, we annotated cells using a pseudo-bulk approach (Figure 3B). We noted that Th1/Th17 cells were the dominant population among all arthritis groups, having the highest proportion in JIA-ERA patients (24% out of all annotated cells). Meanwhile, the proportion of Treg cells in JIA-ERA patients was the lowest among all arthritis groups (only 4% out of all annotated cells). Such imbalance between the pro-inflammatory Th1/Th17 and anti-inflammatory Treg populations is believed to be the cause of inflammation and one of the driving factors in the pathogenesis of JIA-ERA [39]. Interestingly, the different disease groups displayed similar distributions of pseudo-bulk cell types. Although a small sample size could partially explain this finding, nevertheless we believe that different arthritic disease groups might converge to a similar inflammatory joint cell distribution, but with distinct gene expression patterns.

**Figure 3.**
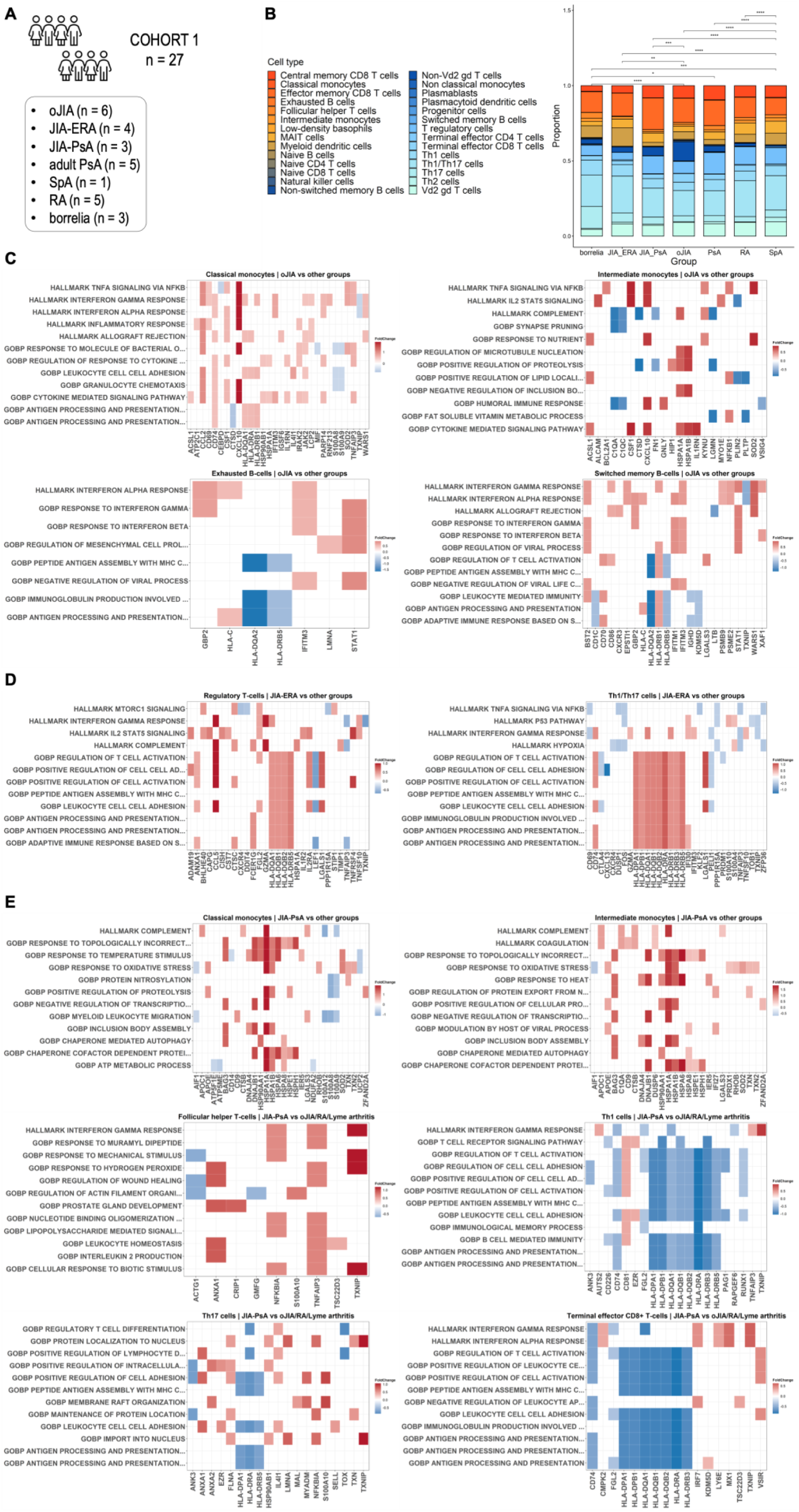
Single-cell phenotype mapping. (A) The cohort included 27 patients having 7 different types of arthritis; (B) Stacked bar plot of the identified cell types, statistical significance was analysed by Wilcoxon test with Benjamini-Hochberg correction (*p ≤ 0.05, **p ≤ 0.01, ***p ≤ 0.001, ****p ≤ 0.0001). Detailed cell counts can be found in Table S5; (C–E) Heatmaps showing the expression of key differentially expressed genes and biological pathways. The most significantly enriched pathways were selected for display. The colour intensity indicates the mean expression level of each gene within a given pathway. Fold change was log2 transformed (p < 0.05).

Regarding gene expression and biological pathway profiles, we observed distinct features in JIA-PsA, JIA-ERA and oJIA compared to other groups (Figure 3C–E). In oJIA patients compared to other groups, CXCL10 was overexpressed in classical and intermediate monocytes. STAT1 was slightly upregulated in exhausted and memory B-cells. Based on the differentially expressed genes in monocytes and B-cells, certain hallmark pathways were enriched such as TNF-α/NF-κB signalling, IL-2/STAT5 signalling, IFN-α and IFN-γ responses, which are relevant to the JAK/STAT pathway. The upregulation of CXCL10 and STAT1, as well as the enriched JAK/STAT-related pathways observed herein suggests the potential use of JAK-inhibitor for the treatment of oJIA. In addition, we noted significant B-cell activation and TNF-α induced activity, the latter thus also supporting the use of TNF-α inhibitors. In Treg, Th1, and Th17 cells of JIA-ERA patients, we noted a significant upregulation of the IFN-γ pathway, CD74 and LGALS1 genes, as well as downregulation of LEF1. Interestingly, we noted in B-cells the upregulation of IGHV7-4.1 which could imply that specific epitopes are causative in JIA-ERA. The presence of CD74 and LGALS1 was observed in various T-cell populations in adult SpA patients [40–42], but not yet for JIA-ERA. CD74 has been found to be an antigen for T-cells and B-cells in SpA, eliciting Th1, Th17, and antibody responses [40,41]. LGALS1 is a member of the galectin protein family that plays a role in regulating cell apoptosis, proliferation, and differentiation, and was found overexpressed in synovial fluid Treg cells of ankylosing spondylitis patients [42]. Thus, our findings further support the concept of the JIA-ERA – SpA continuum. LEF1-deficient Treg cells showed diminished immuno-suppressive capacity and the underexpressed LEF1 was found to result in aberrant T-cell activation and autoimmunity [43]. Its downregulation has been observed upon activation and maturation of Treg cells in JIA patients, suggesting a relatively quiescent/resting phenotype and probably representing Treg cells that only recently migrated into the inflamed joint [44].

Regarding JIA-PsA, there was evidence of upregulated heat-shock protein (HSP) genes in classical and intermediate monocytes. The roles and molecular mechanisms of HSPs that lead to the occurrence and development of psoriasis have been previously reported [45]. Remarkably, we found a substantial upregulation of TXNIP in several T-cell subsets (i.e., Tfh cells, Th1 cells, Th17 cells, terminal effector CD8^+^ T cells), which has previously been reported in psoriatic skin [46] but not yet in psoriatic arthritis. Excitingly, we also note the upregulation of the epithelial migration pathway. Finally, like in oJIA, TNF-α was significantly upregulated in T-cells, thereby supporting the use of TNF-α inhibitors in JIA-PsA.

As T-cells and dendritic cells are known to mediate psoriasis [47], they might also be contributors in the pathogenesis of PsA (Figure S7A). In Tfh cells, Treg cells, and pDC, we observed upregulation of AREG, SOCS1, and ZFP36L2. AREG and SOCS1 were previously identified as psoriasis susceptibility proteins, either in humans or in mice [47]. ZFP36 family members, including ZFP36L1 and ZFP36L2, have been found to regulate inflammatory cytokine production, such as IL-6, CXCL8, and CXCL2, in psoriatic dermal fibroblasts [48]. In NK cells, AREG, SOCS1, and ZFP36L2 were also overexpressed. Interestingly, LAG-3 expression was remarkably low in Treg cells, which agrees with a previous publication that outlined a decline in LAG-3 expressing CD4^+^ T cells in patients with active psoriatic arthritis [49]. Overall JIA-PsA and adult PsA seem to be orchestrated by distinct pathological pathways, which was an unexpected finding. Contrary to patients with PsA, RA patients (Figure S7B) had a high level of LAG3 in their synovial fluid T cells (including Treg cells). A recent report has shown that LAG3 is a phenotypic marker for IL-10 producing Treg cells in PBMCs of RA patients. It was observed that these IL-10 producing LAG3^+^ Treg were less present in PBMCs of RA patients than healthy donors, and were particularly low in frequency in RA patients with higher disease activity, which implied that LAG3^+^ Treg are involved in regulating the pathophysiology of RA [50]. In memory CD8^+^ T cells and Th1 cells, FOS and JUN/JUNB were significantly overexpressed. These proteins are members of the activator protein 1 (AP-1) family and contributors in the AP-1 pathway, which is one of the key pathways that orchestrate the inflammatory responses in RA synovial fibroblasts [51]. Increased levels of FOS and JUN themselves in RA synovium have been reported to correspond with disease severity [52].

### Intercellular communication in adult and juvenile arthritic diseases

After looking into the gene expression of specific cell populations, we continued to investigate how these populations communicate with each other through ligand-receptor interactions. By applying CellChat [53] to our scRNA-seq data, we performed a two-way analysis: first, we analysed each sample individually and then identified the pathways common to each disease across these samples (Figure 4 reflects the most common pathways); the second analysis involved comparing each disease against the others (Figure S8). Details of all identified pathways are documented in Table S6.

**Figure 4.**
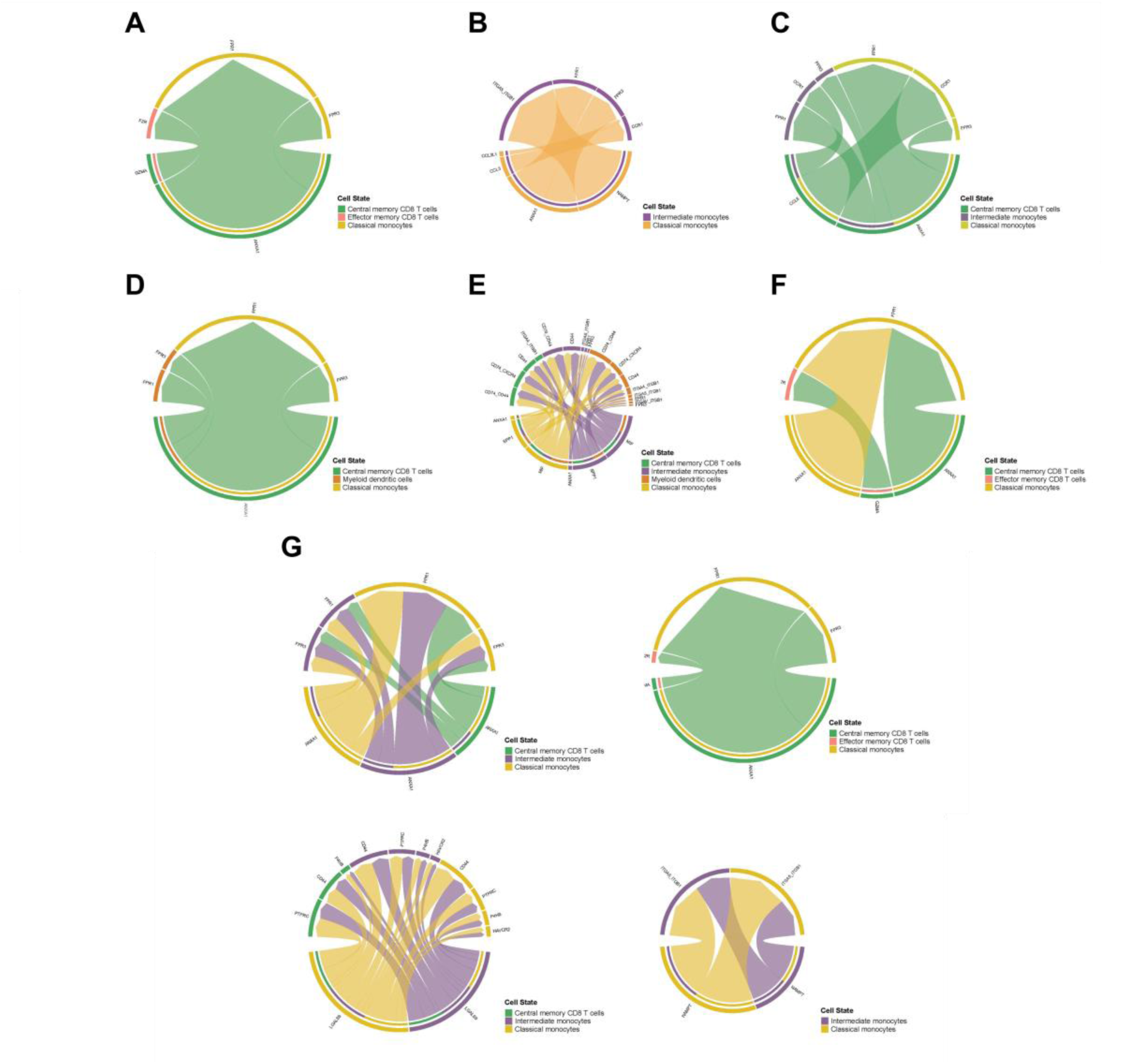
Chord diagrams highlighting the interaction between different cell populations in 7 arthritis groups. (A) Borrelia/Lyme arthritis; (B) JIA-ERA; (C) oJIA; (D) PsA; (E) RA; (F) SpA; (G) JIA-PsA.

As displayed in Figure 4, Annexin signalling seems to be an important pathway present in the intercellular communication of all 7 arthritis groups. In borrelia/Lyme arthritis, CD8^+^ T_CM_ cells communicated mainly with classical monocytes via ANXA1, FPR1, and FPR3 mediators of the Annexin pathway. CD8^+^ T_CM_ cells also interacted with CD8^+^ T_EM_ cells through the PARs pathway with contributions from GZMA and F2R genes. In JIA-ERA, intermediate monocytes communicated with classical monocytes mainly via the Annexin pathway, and also via the Visfatin pathway involving ITGA5, ITGB1, and NAMPT, as well as via the chemokine signalling pathway mediated by CCR1, CCL1, and CCL3L1. In oJIA, CD8^+^ T_CM_ cells interacted with intermediate and classical monocytes by Annexin and chemokine signalling. In adult PsA, CD8^+^ T_CM_ cells communicated with myeloid dendritic cells and classical monocytes via the Annexin pathway. Similarly to borrelia/Lyme arthritis, CD8^+^ T_CM_ cells in SpA patients interacted with CD8^+^ T_EM_ cells and classical monocytes by Annexin and PARs signalling. Synovial fluid of RA patients exhibited strong interaction between CD8^+^ T_CM_ cells, myeloid dendritic cells, intermediate and classical monocytes via not only Annexin signalling, but also MIF and SPP1, which have a crucial role in cytokine activity, immunoregulation, and inflammation. In JIA-PsA, CD8^+^ T_CM_ cells interacted with intermediate monocytes and classical monocytes through not only Annexin signalling, but also Galectin signalling (via CD44, HAVCR2, LGALS9, P4HB, and PTPRC genes). They also communicated with CD8^+^ T_EM_ cells through PARS signalling. Intermediate and classical monocytes communicated with each other by the Visfatin pathway. Furthermore, Figure S8 reveals that several chemokine ligands and receptors, which mediate cellular chemotaxis, were upregulated in 6 out of 7 arthritis groups in our cohort (i.e., borrelia/Lyme arthritis, oJIA, JIA-ERA, JIA-PsA, PsA, and SpA). This suggests the infiltration of immune cells in the joints of these patients.

### Cytokine and chemokine signatures in synovial fluid

To further validate and expand our gene expression findings obtained in our in-house cohort, we next went on to discover disease specific cytokine and chemokine signatures in the synovial fluid from 50 arthritis patients. Although, cytokine analyses in synovial fluid have been done since decades, our approach was strengthened (and made unique) by the simultaneous analysis of the different arthritic-disease groups, thereby minimizing inter-assay variability and allowing appropriate comparison between the different arthritis groups. The 18 oJIA patients had statistically significantly elevated levels of various cytokines (e.g., IFN-γ, IL-10, IL-12p40, IL-13, IL-1RA, IL-2RA, TNF-α, IL-22, and TGF-β) in their synovial fluid compared to the other groups (Figure 5), thereby confirming our GEX-data that showed increased IFNg and TNFa signalling in oJIA. We also confirmed increased IL12 concentrations, which has been observed to increase in oJIA patients during the active period compared to controls [54]. IL-12, or IL-12p70 (70 kDa) is a heterodimeric cytokine composed of a heavy chain p40 (40 kDa) and a light chain p35 (35 kDa) covalently linked to form a biologically active heterodimer. It is produced in the early phase of infection and favours the differentiation of naïve CD4^+^ T-cells into mature Th1 cells [54,55]. The anti-inflammatory cytokine IL-10 was reported to be upregulated in synovial fluid CD4^+^CD25^+^ T cells and CD30^+^ T cells of patients with mild and self-remitting oJIA compared to those with progressive and extended oJIA [34,56]. The upregulation of IL-10 suggests a high level of anti-inflammatory activities which promote immune tolerance. Important chemokines, such as eotaxin (CCL11), Flt1, fractalkine, IP-10, MCP-2, MIP-1α, MIP-1β, and MIP-3β, were abundantly found in synovial fluid of oJIA patients. Eotaxin, a chemokine specific to the anti-inflammatory Th2 cells, was increased in synovial fluid of patients with oJIA, which agrees with the findings from de Jager et al. who reported elevated levels of CCL11 and CCL22 in synovial fluid of oJIA and pJIA patients [57] and suggested that eotaxin may be one of several cytokines that participate in mediating synovial inflammation via Treg recruitment [57,58]. Fractalkine, a chemokine that has been found to be angiogenic in both membrane-bound and free forms, is a mediator of angiogenesis which can foster the infiltration of inflammatory cells into the joints leading to synovial hyperplasia and progressive bone destruction [59]. Pro-inflammatory chemokine MIP-1β and cytokine IFN-γ were previously found relevant in PsA and RA [60–63], but not reported in oJIA.

**Figure 5.**
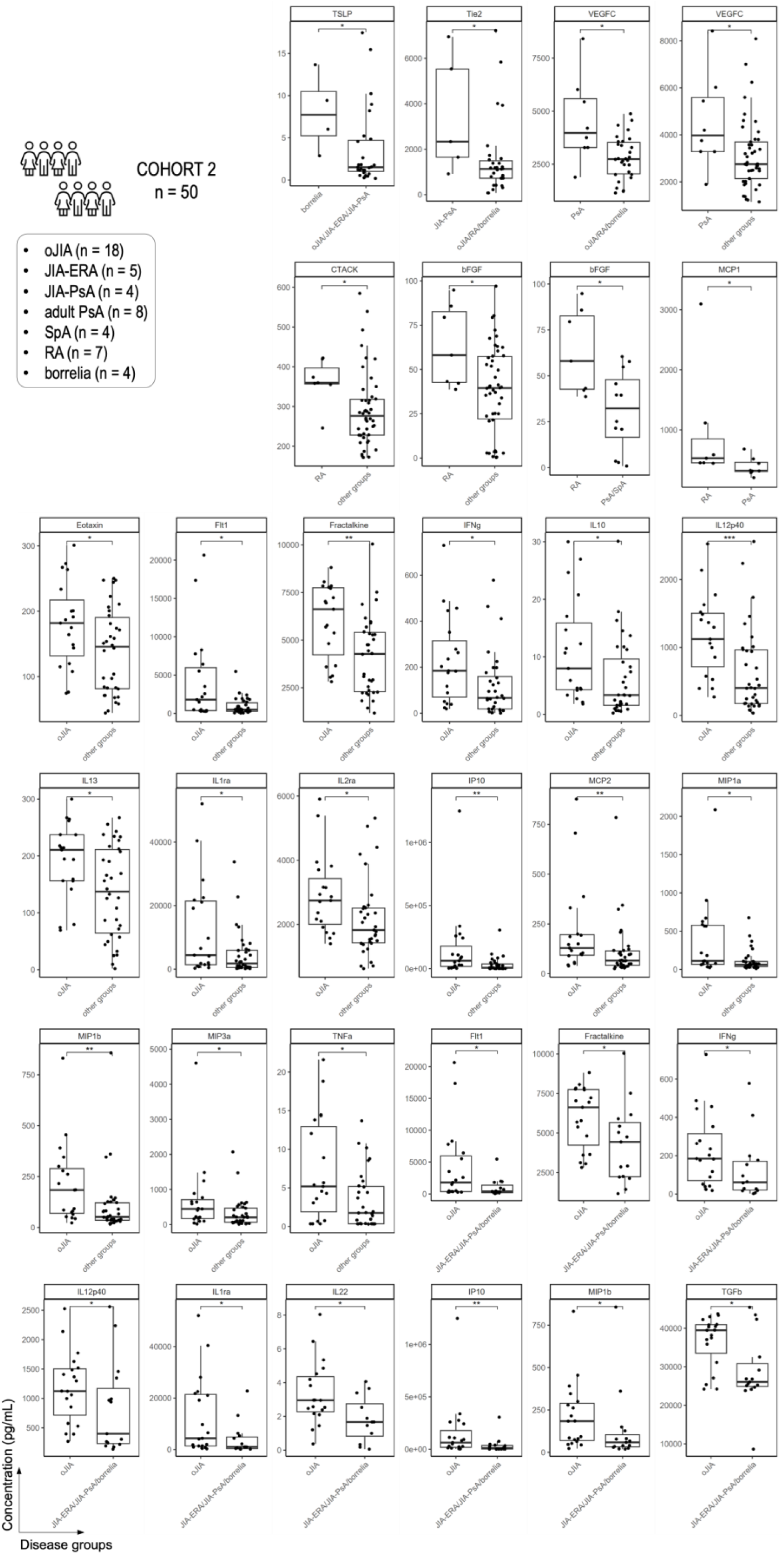
Levels of synovial fluid plasma cytokines and chemokines. Horizontal lines represent median values. Statistical significance analyses were done by Wilcoxon test with Benjamini-Hochberg correction (*p ≤ 0.05, **p ≤ 0.01, ***p ≤ 0.001, ****p ≤ 0.0001). Detailed concentrations can be found in Table S4.

RA patients had significantly higher levels of CTACK, bFGF, and MCP-1 in synovial fluid than other groups. The concentration of synovial fluid bFGF, basic fibroblast growth factor – an angiogenic mediator, was reported to have a positive correlation with white blood cell and neutrophil counts, as well as the severity of radiographic damage, which indicates that bFGF in RA joints may contribute to progressive joint destruction and represent local inflammatory status in affected joints [64]. MCP-1 (or monocyte chemoattractant protein – 1) is a pivotal chemokine for the recruitment of monocytes/macrophages and has previously been found highly expressed in patients suffering from RA [65–67].

Synovial fluid of patients with PsA and JIA-PsA exhibited elevated levels of Tie-2 and VEGF-C. VEGF-C (vascular endothelial growth factor C), a member of the VEGF cytokine family, is another angiogenic mediator. High level of serum VEGF has been proven to be a crucial feature of active psoriatic arthritis, while low level of serum VEGF indicates inactive psoriatic arthritis [68,69]. Synovial fluid of patients with Lyme arthritis (borrelia) had high concentration of TSLP, which is thymic stromal lymphopoietin – a potent pro-inflammatory mediator, which has previously been found to significantly contribute to the immunopathology of RA, particularly T cell activation and joint destruction [70].

Finally, besides our formal statistical analysis of the 7 main arthritic disease groups, we also simultaneously analysed the synovial fluid derived cytokines from other joint conditions including systemic JIA (sJIA, paediatric Still’s disease), pJIA, eoJIA, ACCP^+^ JIA, adult oJIA/pJIA/eoJIA, osteoarthritis, trauma, undifferentiated arthritis, CTLA4-deficiency, IBD, scleroderma (Figure S9). Although the samples obtained from these conditions were too rare to perform statistics, we did note extremely high IL-18 levels in sJIA (only surpassed by the IL-18 level in the inflamed joint of a PAPA patient), thereby confirming the role IL-18 plays in the pathophysiology of Still’s disease, thus not only on a systemic level but also locally in the inflamed joints [71,72]. Our results also seem to confirm that all cytokines (e.g. CXCL10, TNF-α) that were found to be increased in oJIA, were even more increased in pJIA/eoJIA patients, thereby indicating the intuitive idea that the higher severity of pJIA/eoJIA is reflected by higher levels of inflammation (or vice versa). Interestingly, where osteoarthritis seems to be the perfect non-inflammatory “negative control” for almost all cytokines and chemokines tested, we did find higher bFGF levels in osteoarthritis. In fact, the expression of bFGF is highly upregulated in plasma and synovial fluid of primary knee osteoarthritis patients compared to healthy controls, and such elevated bFGF expression was positively correlated to radiographic severity and cartilage degeneration [73,74]. Noteworthy, as noted before bFGF was also found to be higher in RA patients compared to the other arthritis groups.

## DISCUSSION

Modern rheumatology is increasingly evolving in a direction of pathology– and patient–specific diagnostics and therapy (i.e., personalised medicine). Better molecular stratification is thus an absolute necessity to improve patient care. One key transforming factor in rheumatology has been the application of sequencing techniques in assessing the T cell receptor (TCR) and gene expression repertoires in blood cells or synovial fluid cells [75]. Where gene expression analyses (both bulk and single cell) have become state-of-the-art since several years, characterisation of TCR profiles is now rapidly becoming an important complement to understand disease pathogenesis, prognosis, and response to treatment for many diseases. Indeed, over the past few years, several studies have suggested that the TCR repertoire is a valuable indicator of immune monitoring in autoimmune diseases [76,77]. It also led to new treatments, such as when autologous stem cell transplantation was found to aid JIA and dermatomyositis patients via functional renewal and TCR diversification of Treg [78], or when selective deletion of T-cells carrying TRBV9^+^ TCRs using a cytotoxic anti-TRBV9 antibody was tested successfully on an HLA-B27^+^ patient with ankylosing spondylitis [38]. The latter approach, which only eliminates the TCR of interest without systemic immunosuppression, is highly specific and may be applied to other autoimmune disorders. For example, efforts are currently underway to identify common disease-associated TCR motifs in conditions such as type 1 diabetes [79], multiple sclerosis [80], and Crohn’s disease [81].

After applying our novel TRIASSIC tool on our curated inflamed joint TCR database, our study was able to confirm several previously known TCR motifs associated with arthritis disease groups. In addition, we discovered several new TCR motifs that could be associated with specific arthritis groups, such as the TRBV28-RG motif that we linked to oJIA. These motifs could be validated within our independent cohort, indicating that they are highly likely to be disease associated. These are therefore promising candidates for further study into these arthritis groups and may hold promise for therapeutic potential if future research can show that they are driving the disease.

We believe that our TCR data analysis convincingly shows that JIA-ERA and HLA-B27+ SpA target similar epitopes. Moreover, our single cell gene expression analysis further confirmed this statement by showing an upregulation of LGALS1 and CD74 in JIA-ERA patients, a finding that was also previously reported for SpA patients. Interestingly, JIA-ERA patients showed an upregulation of IGHV7-4.1, which could indicate that in HLA-B27+ there isn’t only a sharing of T-cell epitopes but potentially B-cell epitope targets too. Although PsA and SpA (and the juvenile counterparts) are considered to belong to the same disease spectrum, our TCR and gene expression results don’t seem to confirm this, although HLA-B27+ PsA seems to behave in a similar way (at least from the TCR perspective) as SpA.

Regarding oJIA patients both transcriptomics and proteomics data showed various inflammatory pathways upregulated, including interferon type 1 and 2 and TNF-α. Given that in many countries the use of JAK-STAT inhibitors and TNF-α inhibitors are not available for oJIA patients, this finding could be an additional argument in favour of these drugs.

From our extensive analysis of cell-cell interaction based on the scRNA-seq data, Annexin A1 was revealed to be the most prominent pathway in both adult and juvenile arthritis. In an arthritis mouse model, all elements of the Annexin A1 pathway could be detected in naïve joints, with augmentation during the active phase [82]. Our data also highlighted the infiltration of immune cells in the joints as various chemokine ligands and receptors, which mediate cellular chemotaxis, were found upregulated in the synovial fluid of most patients in our cohort (i.e., patients with borrelia/Lyme arthritis, oJIA, JIA-ERA, JIA-PsA, PsA, and SpA) but not RA patients (Figure S8). CD8^+^ T_CM_ cells emerged as an important part of intercellular communication as they engaged with several populations such as CD8^+^ T_EM_ cells, dendritic cells, and monocytes. This emphasizes the interaction between innate and adaptive immune cells in arthritis.

Although our work is robust thanks to the large TCR database (compiled from different studies and experimental approaches), independent in-house validation cohort, the simultaneous analysis of all major chronic arthritis disease groups (thereby minimizing inter-assay and inter-paper biases) and the development of novel computational tools, several limitations should be emphasized. First, we must acknowledge that our validation cohort is modest in size and includes patients that were treated with disease modifying antirheumatic drugs. Another limitation is the imbalance in the sample size between different arthritis groups with oJIA patients having the highest proportion (36%) in the validation cohort, whereas only 8% are JIA-PsA, SpA, or LA patients. The small sample size of certain arthritis groups might be insufficient in accounting for the disease heterogeneity and variability. Despite this limitation, this study generated statistically significant results that can inspire future research endeavours including larger study groups.

In conclusion, we showed that despite the clinical similar presentation of an inflamed joint and despite certain biological commonalities, different chronic arthritic diseases are regulated by different pathophysiological mechanisms that can be discovered when examining the synovial fluid derived TCR, gene expression and cytokine signatures. These findings could lead to the development of novel arthritis diagnostic biomarkers and therapeutics, thereby further leading rheumatology to the era of personalised medicine.

## PATIENTS AND METHODS

### Compilation of publicly available TCR datasets

A database containing TCR sequences derived from synovial fluid was assembled from public literature through an extensive search on Pubmed. Studies were searched using the keywords “Synov* TCR” and selected if TCR sequencing was performed on human synovial fluid in the context of arthritist. TCR sequencing data was extracted from public repositories, including ImmuneAccess and SRA; or requested from the original authors. If called clonotype tables were available (e.g. ImmuneAccess studies), those were used. Alternatively, if the original raw reads in FastQ were available, they were reprocessed using MiXCR version 4.7.0, with the default presets corresponding to library preparation method used in the study. For 10X single-cell data, the standard cellranger filtered_contig_annot file was used as input to extract the clonotypes for the database.

To postprocess the clonotype tables, we developed a unified filtering and processing pipeline to parse repertoires in an identical manner. This included the following steps: 1) mapping V-genes and J-genes to IMGT notation, 2) removal of the unproductive TCR sequences, 3) removal of unproductive TCRs based on an invalid junction or junction regions longer than 30 amino acids, 4) trimming of the nucleotide sequence to the CDR3 region, 5) removal of duplicate clonotypes based on shared nucleotide sequence, 6) processing of the metadata for each repertoire in a common format.

### Patient recruitment

50 patients were recruited at the Divisions of (Paediatric) Rheumatology of Antwerp University Hospital, Department of Orthopedics of Antwerp University Hospital, Divisions of adult and Paediatric Rheumatology of Ziekenhuis Aan de Stroom (ZAS), Divisions of Paediatric Rheumatology of Brussels University Hospital (VUB) and Division of Paediatric Rheumatology Ghent University Hospital. Among these patients, 18 had oJIA, 5 had JIA-ERA, 4 had JIA-PsA, 8 adults had PsA, 7 adults had RA, 4 adults had SpA, and 4 children had Lyme arthritis (LA). Synovial fluid was obtained from the patients by therapeutic aspiration of affected joints. We note that in this study prior treatment was not an exclusion criterium. A clinical overview of patients is presented in Table 1. A list of detailed diagnoses and concurrent treatment can be found in Table S3. This non-commercial academic study was approved by the IRBs from the Antwerp University Hospital/University of Antwerp (IRB numbers 15/43/448, 18/16/215, and 19/30/358) and all local committees from each participating hospital. Written informed consent was collected from the patients (or assent depending on age) and their parents.

### Synovial fluid processing and cryopreservation

Synovial fluid was first treated with hyalorunidase (Sigma Aldrich) at 20 U/mL for 30 minutes at 37°C to break down hyaluronic acid. Synovial fluid mononuclear cells (SFMC) and plasma were isolated using the standard Ficoll density gradient centrifugation procedure. SFMC were cryopreserved in liquid nitrogen and plasma was stored at –20°C.

### Quantification of synovial fluid plasma cytokines and chemokines

For cohort 1 (n = 50), a broad panel of cytokines and chemokines in synovial fluid plasma was quantified via multiplex ELISA (Meso Scale Discovery): BDNF, bFGF, CRP, CTACK, Eotaxin, EPO, Flt1, Fractalkine, GCSF, IFN-γ, IL-10, IL-12p40, IL-12p70, IL-13, IL-15, IL-16, IL-17A, IL18, IL-1RA, IL-1β, IL-22, IL-2RA, IL-4, IL-5, IL-6, IL-8, IL-9, IP-10, MCP-1, MCP-2, MCP-3, M-CSF, MIP-1α, MIP-1β, MIP-3α, Plgf, SAA, sICAM-1, sVCAM-1, TGF-β, Tie2, TNF-α, TSLP, VEGF, VEGF-C, VEGFD. Data were analysed using the MSD Discovery Workbench programme (Meso Scale Discovery). Statistical significance analyses were done in R software using Wilcoxon test with Benjamini-Hochberg correction (*p ≤ 0.05, **p ≤ 0.01, ***p ≤ 0.001, ****p ≤ 0.0001).

### Single-cell RNA and TCR sequencing

For cohort 2 (n = 27), the cryopreserved SFMC were thawed, then dead cells were removed using the Dead Cell Removal Kit (Miltenyi Biotec). The remaining live cells were then stained with hashtag antibodies (BioLegend) for multiplexing, two samples with distinct hashtags would be pooled together. Hashtag-stained cells and barcoded gel beads were loaded onto Chromium Next GEM Chip K (10X Genomics) and partitioned in oil droplets using the Chromium Controller (10X Genomics). The targeted number of recovered cells was 10,000 per multiplexed sample. All subsequent steps were performed based on the Chromium Next GEM Single Cell 5’ Dual Index with Feature Barcode v2 protocol (10X Genomics). Libraries were sequenced on the Illumina NovaSeq 6000 with the sequencing depths as follows: 30,000 read pairs per cell for gene expression libraries; 5,000 read pairs per cell for surface protein libraries; and 5,000 read pairs per cell for V(D)J libraries.

The Cell Ranger *multi* pipeline v7.1.0 was used to demultiplex the hashed 5’ gene expression FASTQ libraries and assign reads to individual samples. Next, the resulting sample-specific BAM files were converted back to FASTQ files, using the Cell Ranger *bamtofastq* function. A final run of the *multi* pipeline was performed to map the sample-specific gene expression FASTQ library reads to the GRCh38 reference genome while simultaneously align the reads from the V(D)J library to the V(D)J-compatible GRCh38 7.0.0 reference set and generate the antibody feature barcode count matrix. The resulting count matrices were analysed using the R package *Seurat* [83]. The data of individual samples were integrated using Seurat’s *merge* command, and subsequently inspected to confirm for a lack of batch effects. To remove multiplets and other low-quality cells, all cells expressing more than 8000 genes, above 20% mitochondrial genes, and 40% ribosomal genes were removed. The data was then log-transformed, scaled, and centred. Next, the most variable genes were identified and used as input for a principal component analysis. The top principal components were then used for clustering, dimension reduction, and visualization using the Uniform Manifold Approximation and Projection (UMAP) method [84]. The resulting cell clusters will be annotated using an *ensemble* method of reference-based mapping based on the *SingleR* package. Differentially expressed genes (DEGs) were identified using the *FindMarkers* function in Seurat with the default Wilcoxon Rank Sum test. A log_2_ fold change threshold of 0.25 was applied, and only genes with a Benjamini-Hochberg adjusted p-value < 0.05 were considered for further analyses. The identified DEG will be used for gene set enrichment analysis (GSEA) using the *clusterProfiler* package. Pathway annotation will be performed based on the Gene Ontology Biological Process and Hallmark gene sets from MSigDB [85].

### Bulk TCR sequencing

For cohort 3 (n = 17), synovial fluid was used for bulk TCR sequencing. This cohort included 6 oJIA patients, 3 JIA-ERA patients, 2 JIA-PsA patients, 2 adult PsA patients, 3 borrelia patients, and 1 RA patient. SFMC were first sorted into three distinct subsets: regulatory T cells (CD3^+^CD4^+^CD25^+^CD127^−^), helper T cells (CD3^+^CD4^+^CD8^−^, excluding the regulatory T cells), and cytotoxic T cells (CD3^+^CD4^−^CD8^+^). Sorting was performed on the FACSAria II instrument (BD Biosciences) at the Flow Cyometry Core Facility (University of Antwerp). Up to 50.000 cells were sorted for each patient per subset. Sorted cells were stored in DNA/RNA Shield (Zymo Research) at –80°C until use. The stabilised cells were then processed using the Quick-RNA Microprep Kit (Zymo Research) for RNA extraction. TCR libraries of the three T cell populations were prepared separately from the eluted RNA using the QIAseq Immune Repertoire RNA Library Kit (Qiagen). All TCR alpha, beta, gamma, and delta chains were amplified and sequenced. Sequencing was done on a NextSeq 500 (Illumina) at the Centre for Medical Genetics (University of Antwerp) as described in one of our previous studies [86].

### TCR immunoinformatics

#### Convergent clonotypes

Identification of the convergent clonotypes was based on the workflow as depicted in Figure S1. 1) Removal of the duplicate TCRs, retaining only those that are the result of an independent recombination event. Hence, clonotypes were defined as unique combinations of a V gene + CDR3 NT + individual. 2) Similar TCR pairs were identified using the clustcrdist tool (Valkiers et al., in preparation), where similarity is defined as those with a TCRdist distance score of less than 12.5 (Mayer-Blackwell et al). 3) For each TCR in the dataset, the number of similar yet independently arising TCRs within the database was counted. These coincidences could occur within one patient, to identify convergent clonotypes within a single individual, but also across repertoires, to identify convergent clonotypes within a group. 4) A TCR background was then defined for each query TCR as its 2048 most similar TCRs based on the same TCRdist metric. We assume that this set of TCRs is large enough to span many epitopes, yet small enough to control for technical bias, e.g. differences in V-gene abundances in the control repertoires against the database. 5) Significantly convergent TCRs were then identified by enrichment in either the database or controls, based on a two-tailed fisher exact test. 6) For each query TCR, the convergence score was computed, describing the log fold change in the number of neighbors in arthritis vs background TCRs:

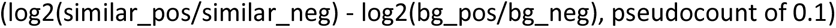

#### Clustering significantly convergent TCRs into SFCCs

Given that significantly convergent TCRs will occur together, we performed clustering using ClusTCR. Given that the set of unique convergent TCRs is more clonal than normal TCR repertoires, we modified the ClusTCR parameters (mcl_inflation=1.5, mcl_expansion=2) to improve cluster purity. Cluster size was defined as the number of unique convergent TCRs from this cluster. The convergence score for a cluster was defined as the mean convergence score of its TCRs. Furthermore, we computed the publicity as the fraction of arthritis individuals that have at least one non-background TCR in the cluster.

#### Mapping the SFCC space through co-occurrence

### Publicity analysis

Given that TCR convergence was computed against an ‘epitope-naive’ simulated background based on shared V/J usage, the resulting SFCCs could simply be MAIT cells or common epitopes that are not necessarily unique to SF. Hence, we assessed the presence of each significantly convergent clonotype in a compiled dataset of non-arthritis PB TCR-seq results (PB control data). This control cohort comprises the studies in Table S1. We then determined the extent of ‘uniqueness’ of a SFCC within SF and arthritis by following two metrics: (i) Hit rate lfc: the log fold change of the frequency of the TCRs in the arthritis database versus the PB control data, (ii) N repertoires lfc: log fold change of the number of repertoires in which the TCR occurs in the arthritis database versus the PB control data.

#### TCR HLA association and epitope specificity prediction

To map the HLA-specificity to individual TCR sequences, we used the list of HLA-associated TCR as described in [87]. This list was queried based on identical V/J/CDR3 usage. Only exact matches were retained as HLA-associated. If a TCR was associated with more than one HLA, only the most significant association was retained.

The epitope specificity of TCR sequences was annotated using DETECT version 1.0. developed by ImmuneWatch BV, available at: https://www.immunewatch.com/detect. The recommended settings were applied, where a score higher than 0.23 was considered as a reliable epitope annotation.

### Cell-Cell interaction meta-analysis

We employed the CellChat R package (v2.1.0), a robust toolkit for analyzing and visualizing cell-cell communication networks from scRNA-seq data, to investigate cell-cell communication in inflamed joints of arthritis patients [53]. CellChat allowed us to infer signaling pathways and ligand-receptor interactions occurring between various cell types, providing insights into disease-specific communication networks. To enhance the robustness of our findings, we performed a meta-analysis across all samples, combining individual CellChat results and applying statistical tests, including Fisher’s combined probability test. This approach helped identify consistent and potentially critical cell-cell interactions implicated in arthritic disease pathogenesis. For network construction, we used the createCellChat function and computed communication probabilities between cell types with computeCommunProb, filtering relevant communication pathways. Additionally, we compared interaction networks across groups (e.g., using the compareInteractions and rankNet functions) to identify disease-specific patterns. Visualization functions, such as netVisual_circle and netVisual_heatmap, highlighted differences in cell-cell communication. This comprehensive analysis allowed us to uncover unique signaling patterns and interactions that may serve as therapeutic targets.

## ACKNOWLEDGEMENTS

We appreciate the participation of all patients and their families in this study. We are grateful to all unmentioned clinicians, nurses, and lab colleagues. We thank Dr George Elias for helping us with the initial sample processing and storage.

