## Supplementary Materials for "Generation of a T cell receptor, cytokine and cell repertoire synovial fluid atlas to define commonalities and dissimilarities between arthritic diseases through systems immunology approaches"

### SUPPLEMENTARY TABLES

**Table S1.** Studies used in the synovial fluid TCR sequencing database.

| Study ID | Sequencing Method | N repertoires | N patients | Diseases | Depth per patient (median) | **Reference** |
|---|---|---|---|---|---|---|
| Dunlap_2024 | 10X | 13 | 13 | RA | 1102 | |
| Henderson_2014 | ImmunoSEQ | 26 | 13 | PsA, oJIA, LA | 19683 | |
| Komech_2022 | ImmunoSEQ | 64 | 24 | PsA, SpA | 14585 | |
| Maschmeyer_2021 | 10X | 21 | 7 | oJIA, eoJIA | 4117 | |
| Mijnheer_2023 | 5-RACE | 36 | 7 | oJIA, eoJIA, pJIA | 19467 | |
| Penkava_2020 | 10X | 5 | 5 | PsA | 7138 | |
| Rossetti_2017 | ImmunoSEQ | 11 | 11 | oJIA/pJIA | 2100 | |
| Savola_2017 | ImmunoSEQ | 3 | 2 | RA | 2351,5 | |
| Spreafico_2016 | ImmunoSEQ | 6 | 6 | oJIA/pJIA | 4546,5 | |
| Steel_2020 | ImmunoSEQ | 9 | 3 | PsA | 4399 | |
| Venken_2024 | 10X | 14 | 14 | RA | 118,5 | |
| Durham_2024 | 10X | 7 | 6 | PsA, RA | 5152,5 | |
| Morbach_2024 | iRepertoire | 15 | 11 | LA | 7475 | |
| Povoleri_2023 | 10X | 4 | 4 | PsA | 3485,5 | |
| Zheng_2024 | other | 8 | 4 | RA | 2799 | |

**Table S2.** Studies used as control (non-arthritis) PB TCR repertoires. For the ImmunoCODE study, samples from Delmonte_2023 were not included.

| Study ID | N repertoires | N patients | Depth per patient (median) | Reference |
|---|---|---|---|---|
| Vlasova_2023 | 1250 | 1225 | 11499 | https://doi.org/10.1101/2023.11.08.566227 |
| Delmonte_2023 | 778 | 778 | 56917 | https://doi.org/10.1016/j.jaci.2023.12.011 |
| ImmuneCODE | 704 | 704 | 124650 | https://clients.adaptivebiotech.com/pub/covid-2020 |
| Emerson_2017 | 666 | 666 | 146498 | https://doi.org/10.1038/ng.3822 |
| Musvosvi_2022 | 276 | 215 | 94813 | https://doi.org/10.21417/MM2022NM |
| Russell_2022 | 150 | 150 | 8965 | https://doi.org/10.7554/eLife.73475 |
| Milighetti_2023 | 182 | 46 | 62839 | https://doi.org/10.1016/j.isci.2023.106937 |

### SUPPLEMENTARY FIGURES

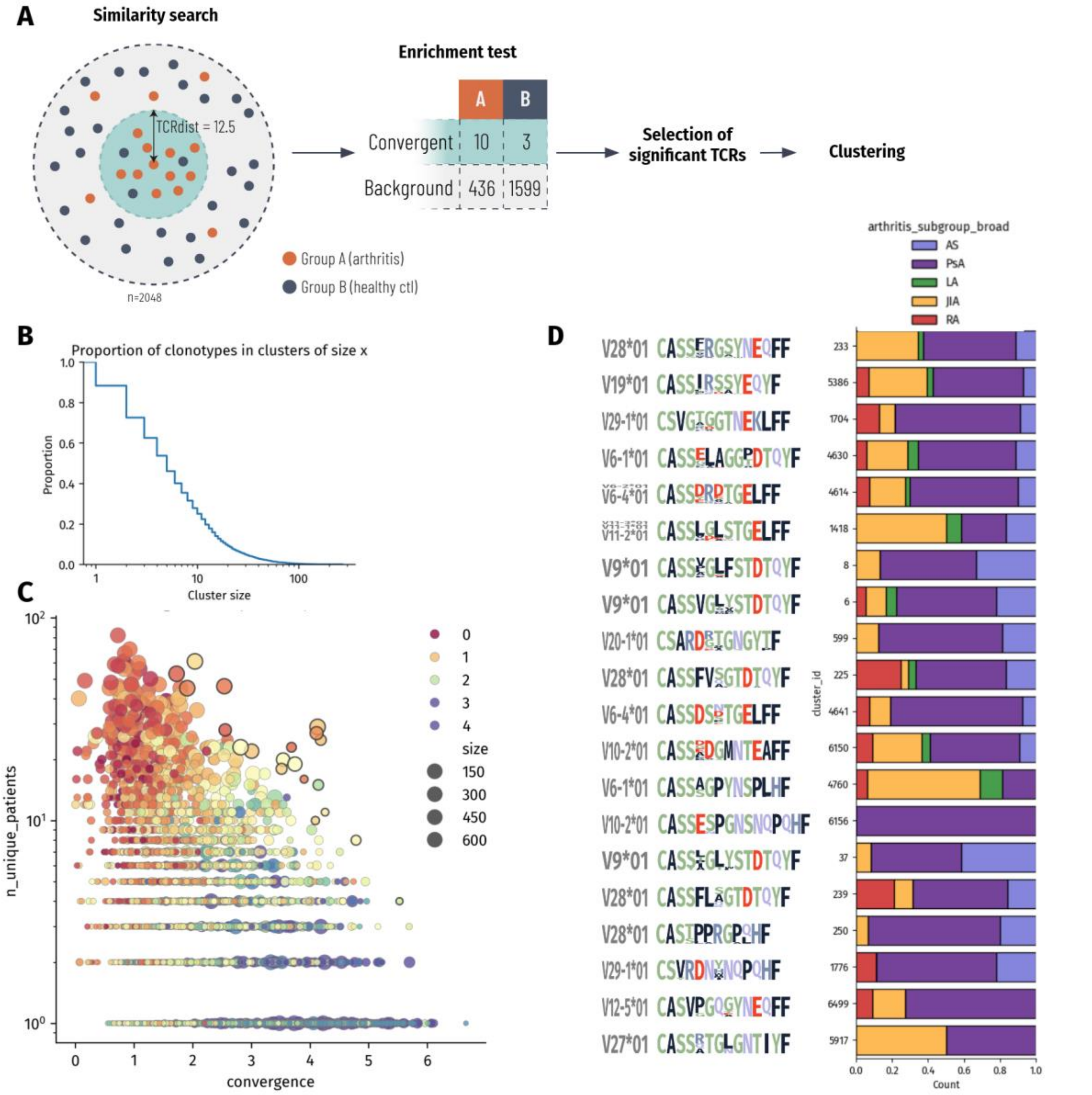


**Figure S1 Convergence analysis overview. (A)** TRIASSIC identifies regions of high convergent recombination in TCR repertoires. We define clonotypes as unique combinations of patient ID, junction nucleotide sequence, V-gene, and J-gene, ensuring each represents an independently generated TCR. For each clonotype, we define its local context as its 2048 most similar neighbors in TCRdist space. Within this neighborhood, we compare the proportion of arthritis-associated versus background clonotypes that are highly similar (TCRdist ≤ 12.5, e.g., 10:3) to their distribution in the non-similar broader neighborhood (e.g., 436:1599). This local analysis approach accounts for sequence similarity and minimizes the impact of global technical variations such as V-gene distribution biases. We assess significance using a one-tailed Fisher exact test, then group significantly convergent TCRs into synovial fluid convergence clusters (SFCCs) using ClusTCR. **(B)** Cumulative distribution of clonotype cluster sizes. The y-axis shows the proportion of clonotypes that belong to clusters of size x or larger, where x is shown as a logarithmic x-axis. **(C)** Relationship between SFCC convergence score and publicity. Each point represents a SFCC, with color indicating its log2 fold enrichment in arthritis SF/ST compared to healthy PB. Black outlines highlight notable SFCCs that show both high convergence and relatively broad sharing across patients. **(D)** Detailed analysis of notable SFCCs identified in (C). Left panel shows V-gene usage and CDR3 sequence patterns. Right panel displays the distribution of these SFCCs across different arthritic diseases.

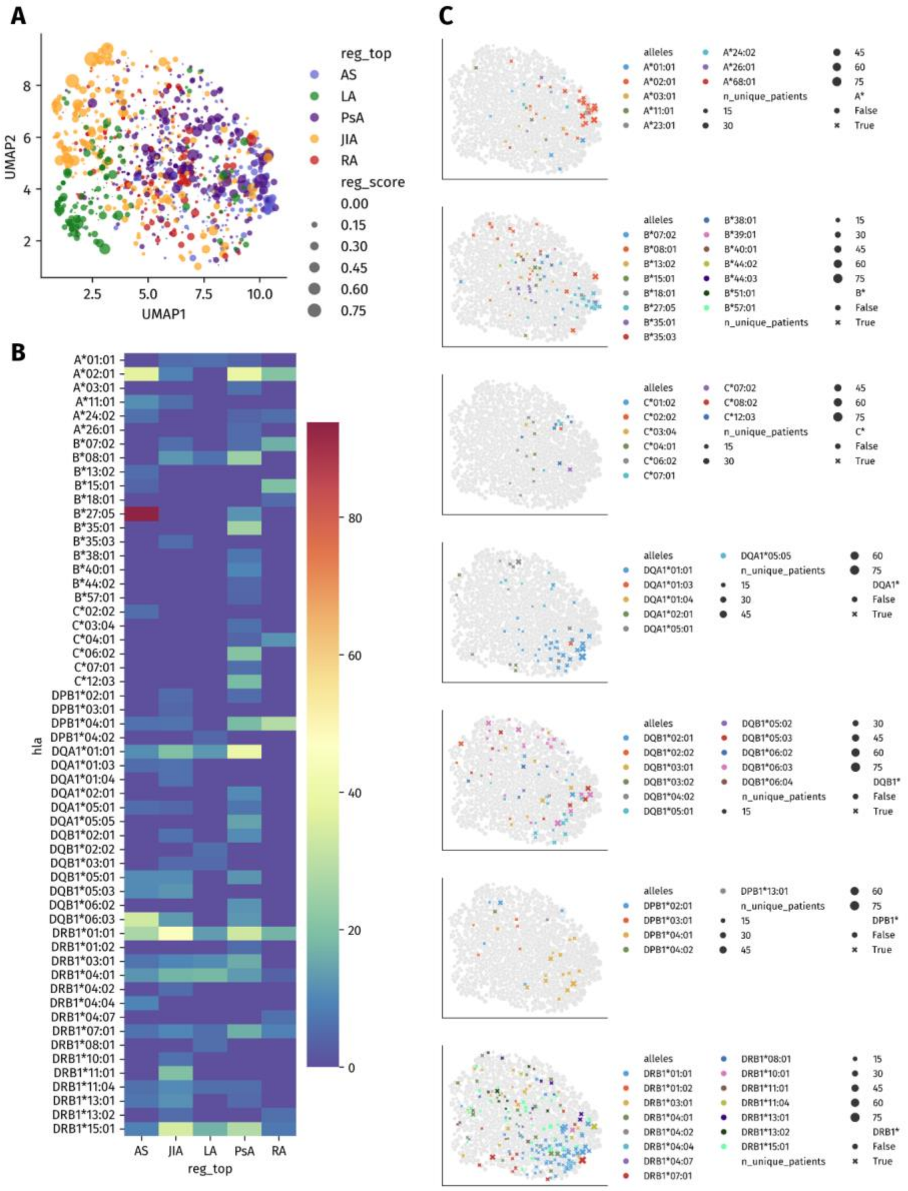


**Figure S2 HLA association of SFCCs.** We investigated the presence of TCRs that are known to be HLA-associated, and their distribution within the SFCCs. **(A)** Co-occurrence mapping reveals relationships among SFCCs using UMAP dimensionality reduction. Each dot represents a SFCC, and their spatial arrangement was determined as the cosine similarity between patient-level vectors (n=130 patients). Proximal points represent SFCCs with similar patient distribution patterns. Dot size represents the strength of association with arthritis disease subgroups determined by multiple logistic regression, while color indicates the diseases with the strongest association. **(B)** Overview of the number of HLA-associated SFCCs per disease-association group. **(C)** Distribution of the HLA-specific SFCCs across the SFCC co-occurrence space. SFCCs specific to the same HLA are frequently located close together. Note that e.g. HLA-B27 associated SFCCs are located in a region with high density of AS-specific SFCCs.

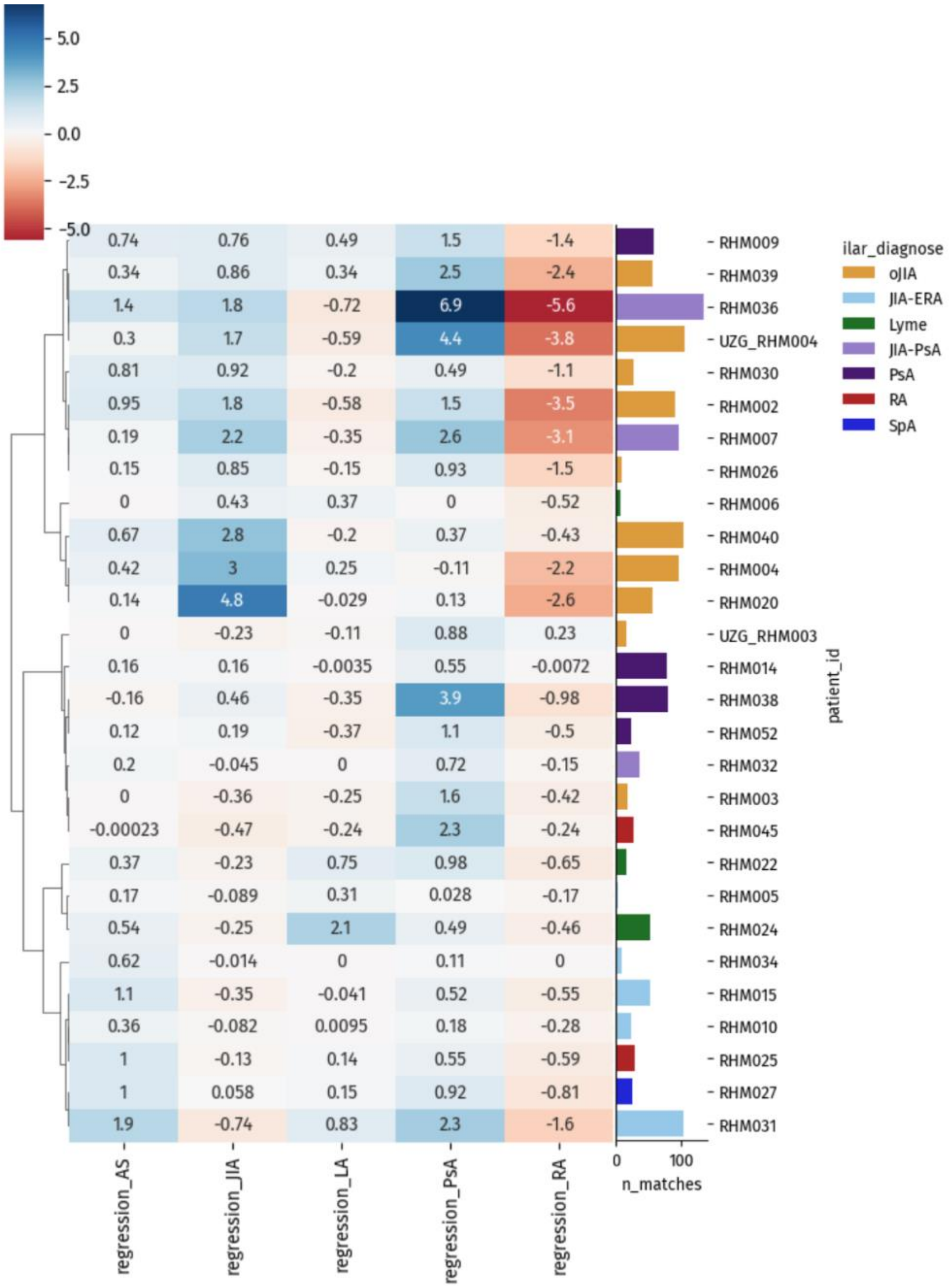


**Figure S3. Unsupervised clustering of in-house arthritis patients based solely on disease-associated SFCCs.** Values indicate the summed regression coefficients for the clonotypes that were also present in the SFCCs, per disease. Repertoires were hierarchically clustered, using cosine similarity to minimize the effect of repertoire size. The histogram to the right shows the number of clonotypes also present within the SFCC database, with the colour representing the disease label. This unsupervised approach does not fully delineate different arthritic diseases. For example, JIA-PsA patients are clustered with the oJIA patients, indicating that these patient groups cannot be easily differentiated based on their TCR-repertoire.

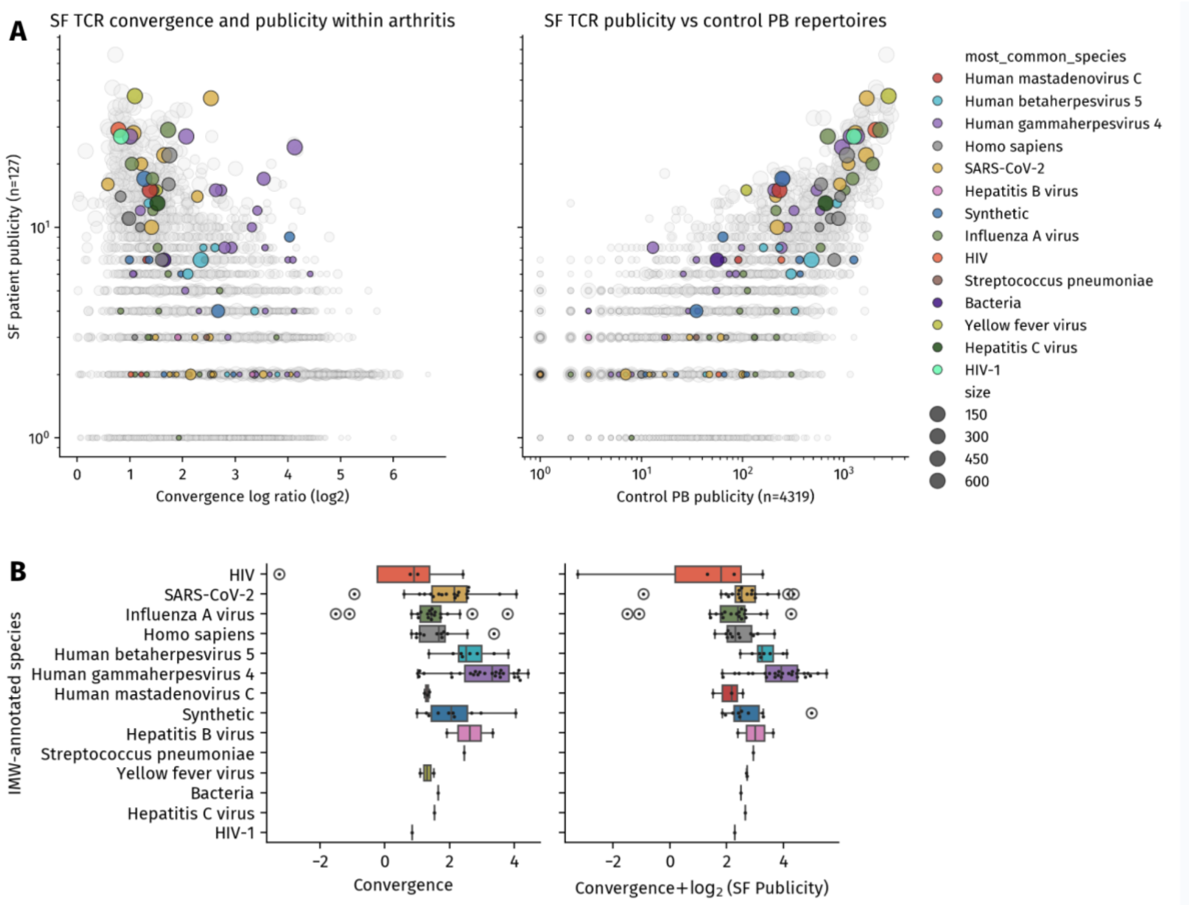


**Figure S4.** SFCCs against Human gammaherpesvirus (Epstein-Barr virus, EBV) show pronounced convergence in SF/ST of arthritis patients. **(A)** Left: Convergence (x-axis) vs number of samples in which SFCC occurs (y-axis). Right: Publicity of viral reactive TCRs in non-arthritis control PB repertoires (x-axis, Table S2). SFCCs are colored based on the specificity of their cognate epitope, as determined by IMW-detect. SFCCs that could not be annotated with high confidence are shown in grey. **(B)** Significantly enriched convergence of SFCCs reactive against EBV vs others (Kruskal-Wallis test p=2.185e-4, MWU for EBV vs rest p=2.812e-6), which is even more apparent when taking publicity of the SFCC into account (Kruskal-Wallis test p= 1.350e-04, MWU for EBV vs rest p=2.584e-07).

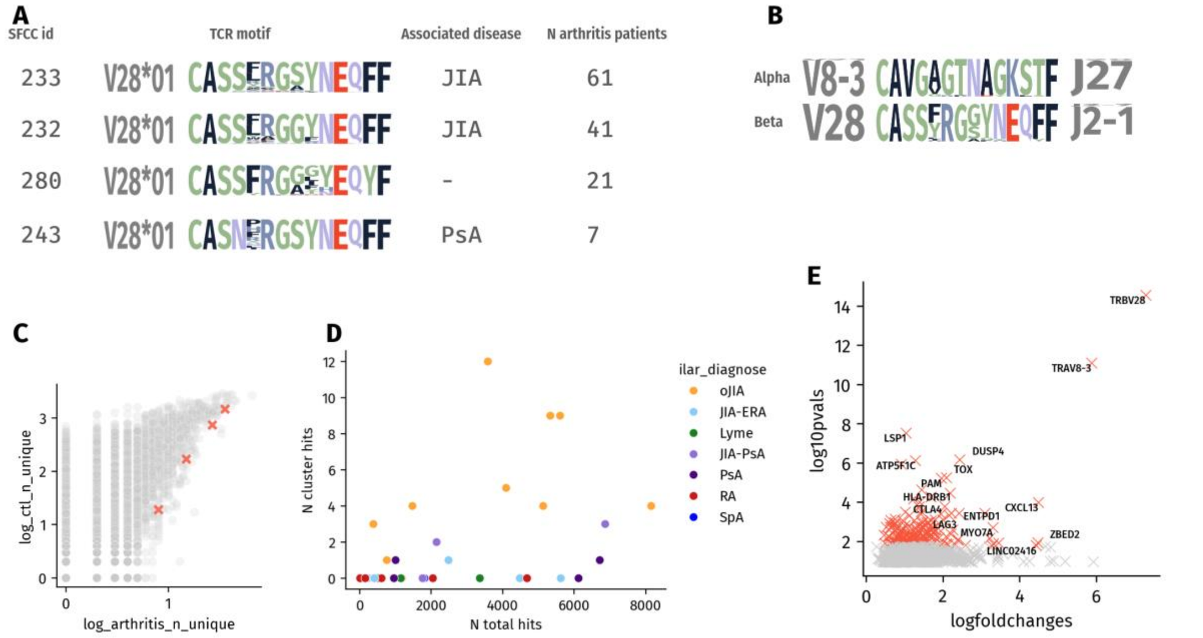


**Figure S5. A convergent TRBV28 TCR with conserved motif, present across four SFCCs, is associated with oJIA.** **(A)** Four widely shared SFCCs with a conserved motif were found in the arthritis SF/ST by TRIASSIC. The largest two SFCCs were predictive for patients with oligo/polyarticular JIA within the database, while the smallest was weakly associated with PsA. **(B)** The TCRs from these SFCCs were also present in our in-house arthritis data. Interestingly, they were paired with a conserved alpha chain, further indicating that they likely recognize the same epitope across patients. **(C)** Prevalence analysis of SFCCs in patient samples. Each point represents one SFCC, with its frequency in arthritis synovial fluid/tissue (n=130) plotted against healthy peripheral blood (n=4319) on log2 scales. A pseudocount of 1 was added to accommodate zero values. **(D)** Independent validation using our in-house dataset confirmed strong enrichment of these SFCC sequences in oligoarticular JIA. The TCR hit frequency was significantly higher in oJIA compared to other diseases (Mann-Whitney U test, p=3.716e-05). **E.** Differential gene expression analysis of T-cells with this specificity (vs other T-cells) revealed a CXCL13-producing T cell signature.

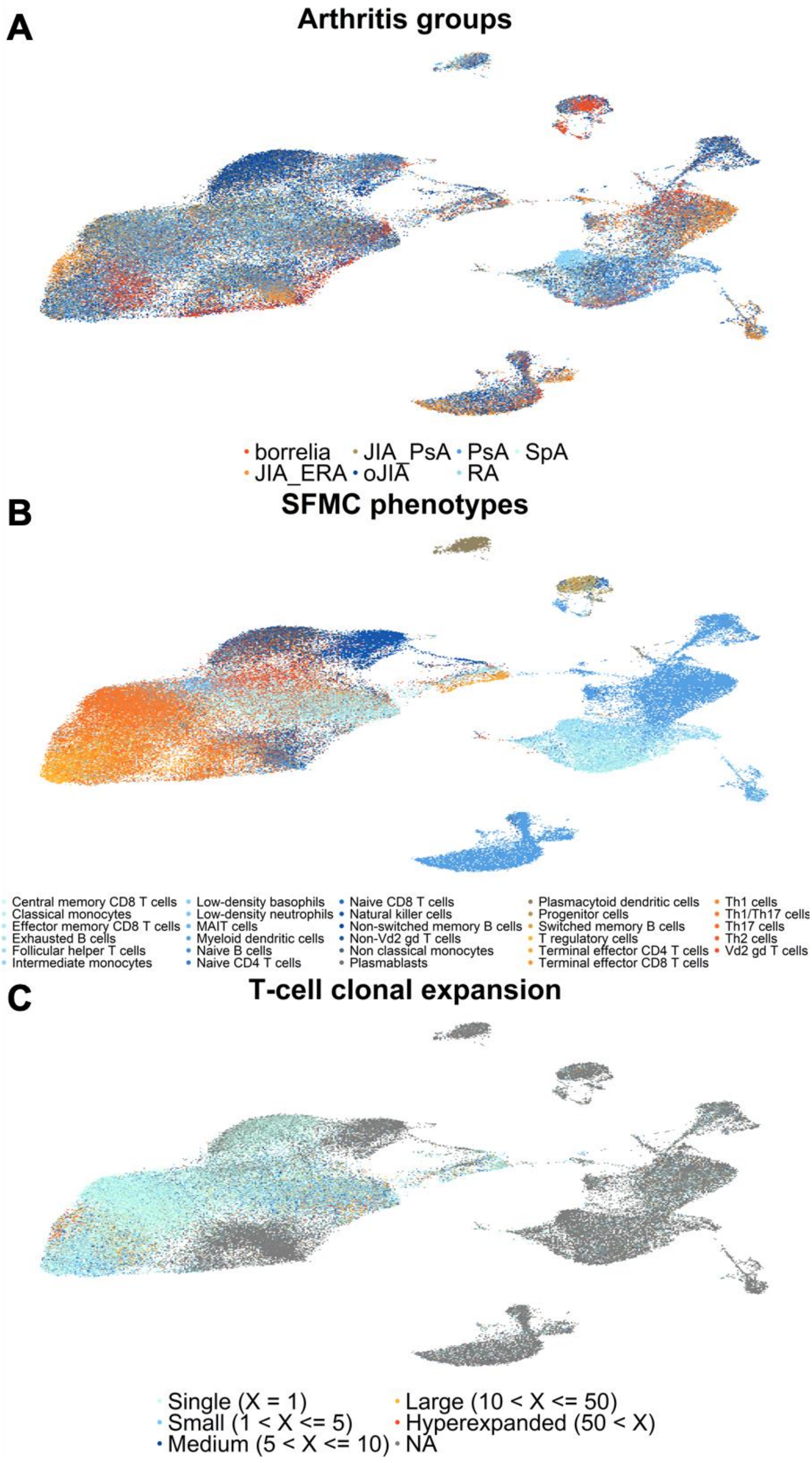


**Figure S6. Single-cell annotation by UMAP. Each dot represents one cell and the colour of each dot represents the group the cell belongs to. (A)** UMAP representation of arthritis groups; **(B)** UMAP representation of gene expression-based phenotypes; **(C)** UMAP representation of T-cell clonal expansion.

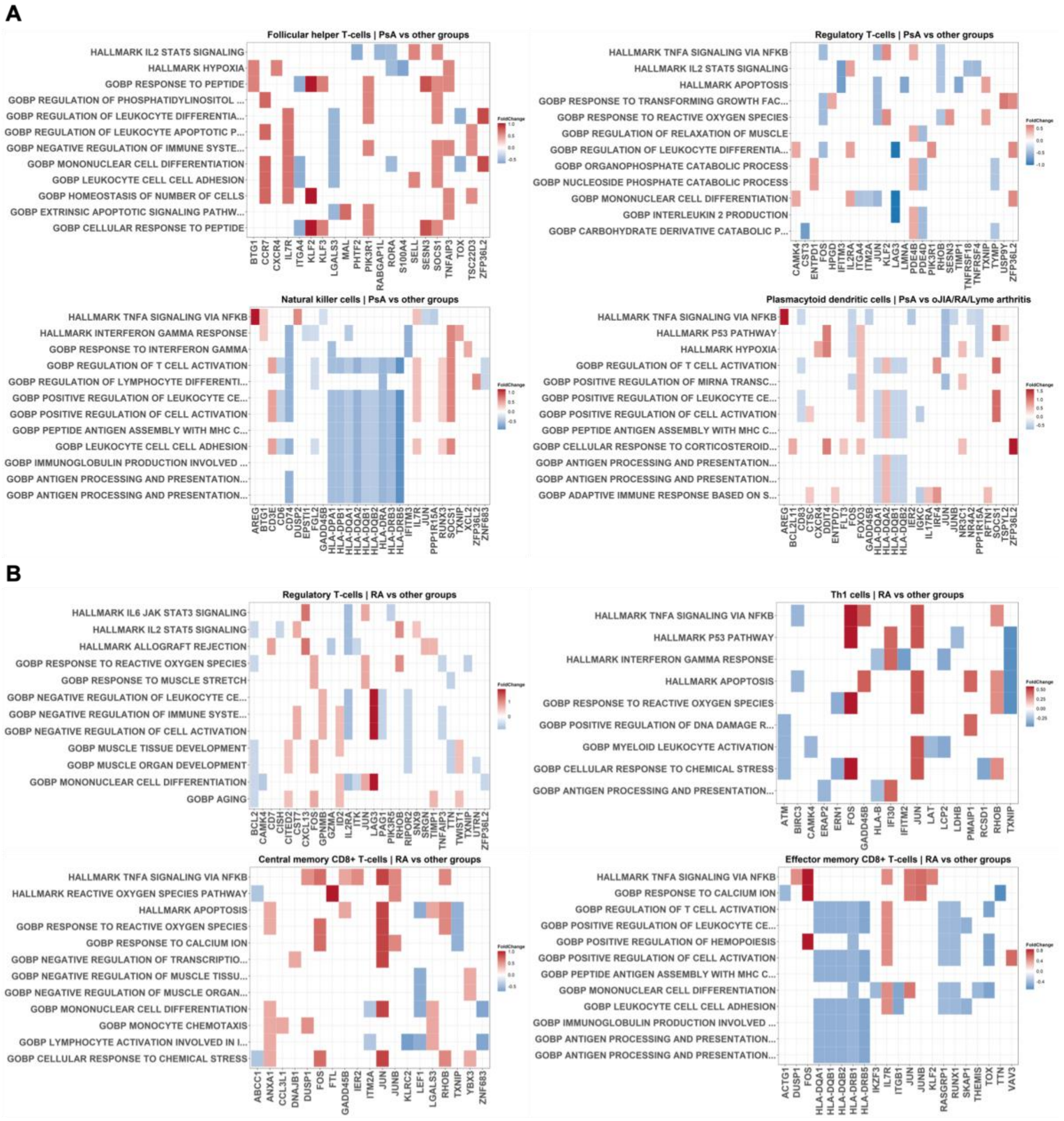


**Figure S7**. **Heatmaps showing the expression of key differentially expressed genes and biological pathways.** The most significantly enriched pathways were selected for display. The colour intensity indicates the mean expression level of each gene within a given pathway. Fold change was log2 transformed ($p < 0.05$). **(A)** Comparison between PsA and other groups; **(B)** Comparison between RA and other groups.

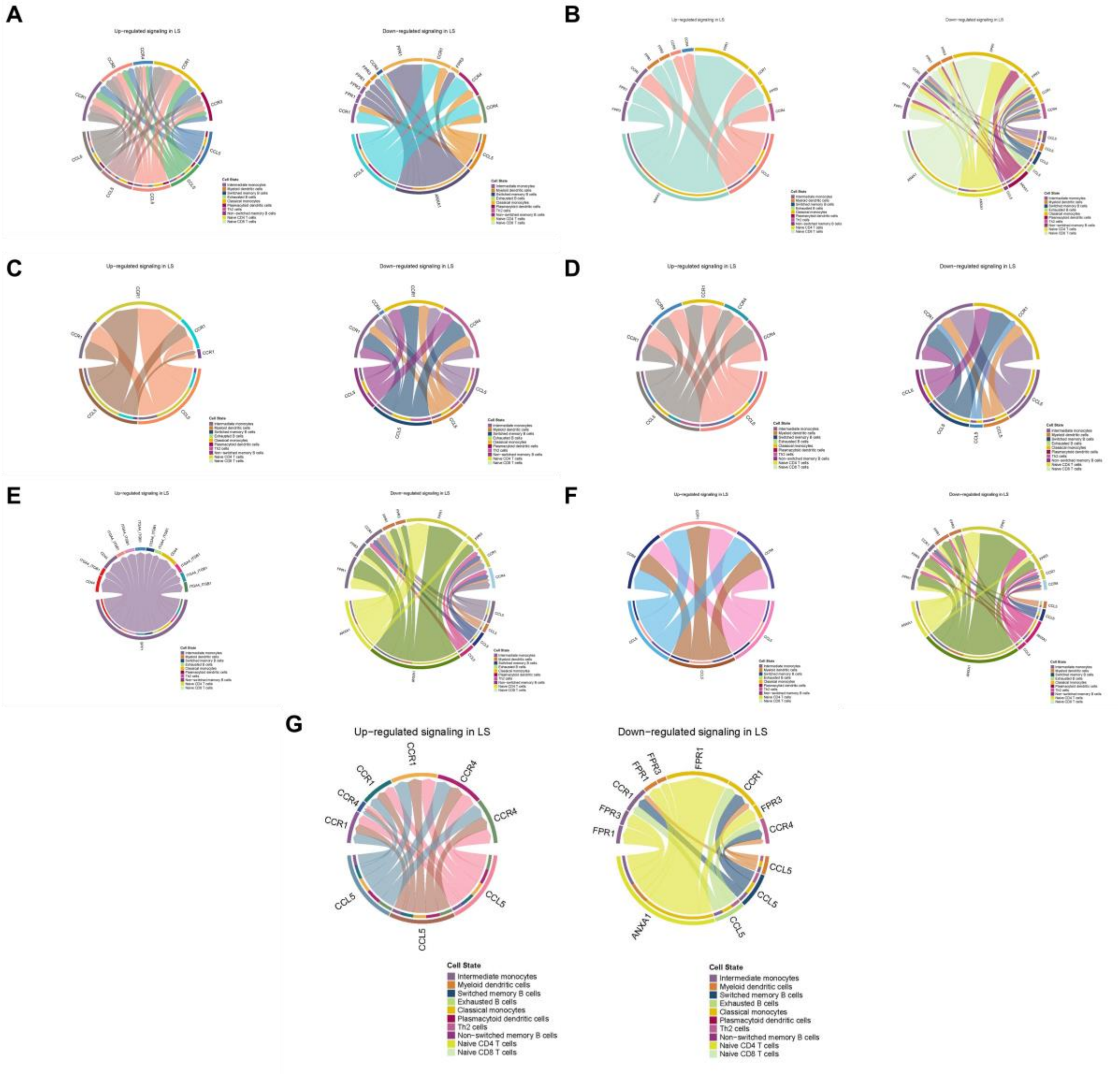


**Figure S8. Chord diagrams comparing the cell-cell interaction between 7 arthritis groups. (A)** Borrelia/Lyme arthritis against the others; **(B)** JIA-PsA against the others; **(C)** oJIA against the others; **(D)** PsA against the others; **(E)** RA against the others; **(F)** SpA against the others; **(G)** JIA-ERA against the others.

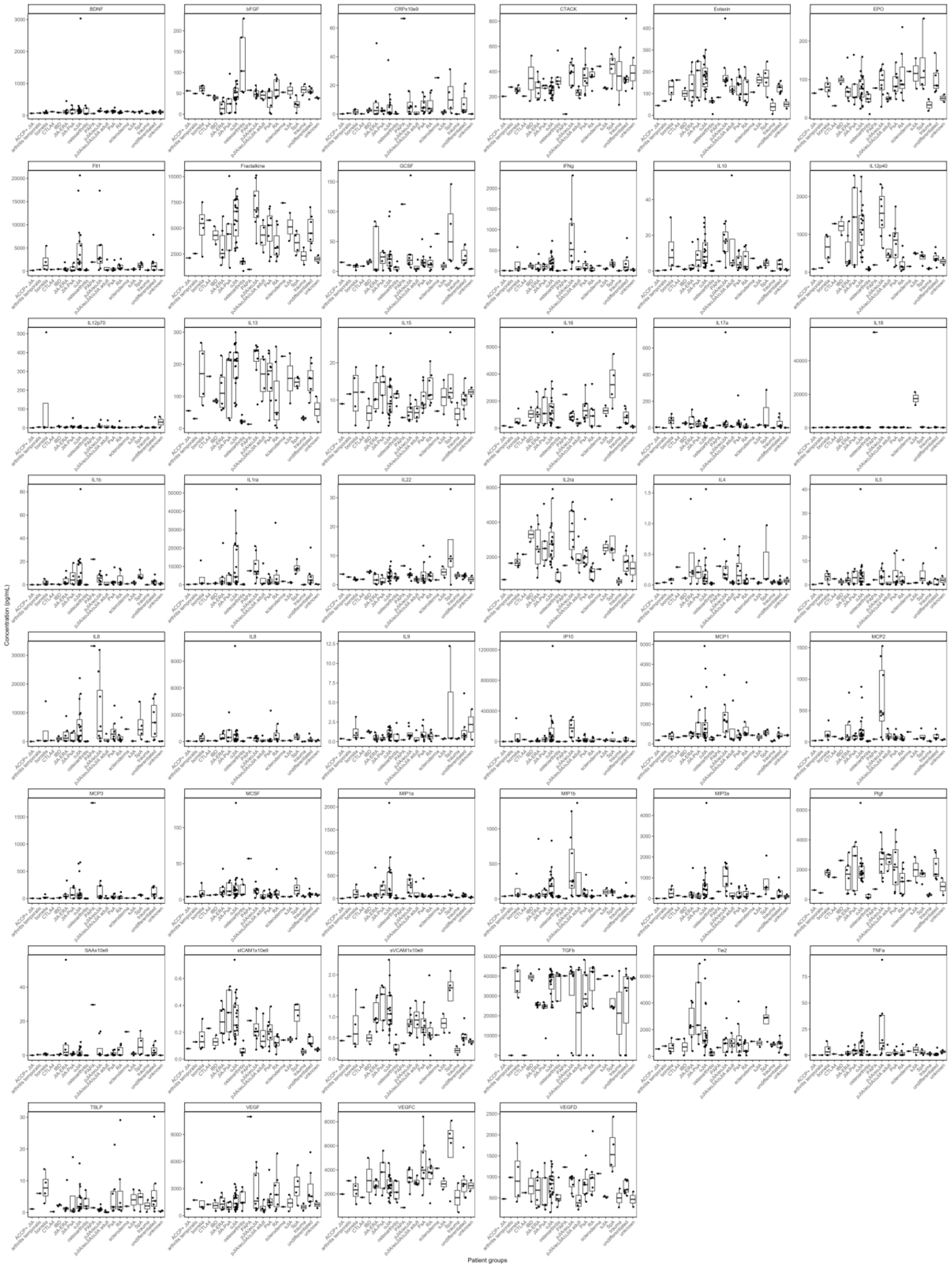


**Figure S9. Levels of synovial fluid plasma cytokines and chemokines.** Horizontal lines represent median values.
